# scCensus: Off-target scRNA-seq reads reveal meaningful biology

**DOI:** 10.1101/2024.01.29.577807

**Authors:** Dongze He, Stephen M. Mount, Rob Patro

## Abstract

Single-cell RNA-sequencing (scRNA-seq) provides unprecedented insights into cellular heterogeneity. Although scRNA-seq reads from most prevalent and popular tagged-end protocols are expected to arise from the 3′ end of polyadenylated RNAs, recent studies have shown that “off-target” reads can constitute a substantial portion of the read population. in this work, we introduced `scCensus`, a comprehensive analysis workflow for systematically evaluating and categorizing off-target reads in scRNA-seq. We applied `scCensus` to seven scRNA-seq datasets. Our analysis of intergenic reads shows that these off-target reads contain information about chromatin structure and can be used to identify similar cells across modalities. Our analysis of antisense reads suggests that these reads can be used to improve gene detection and capture interesting transcriptional activities like antisense transcription. Furthermore, using splice-aware quantification, we find that spliced and unspliced reads provide distinct information about cell clusters and biomarkers, suggesting the utility of integrating signals from reads with different splicing statuses.

Overall, our results suggest that off-target scRNA-seq reads contain underappreciated information about various transcriptional activities. These observations about yet-unexploited information in existing scRNA-seq data will help guide and motivate the community to improve current algorithms and analysis methods, and to develop novel approaches that utilize off-target reads to extend the reach and accuracy of single-cell data analysis pipelines.

**Data Availability:** The scripts for reproducing the results are available at https://github.com/COMBINE-lab/sc-census. Supplementary files can be found at https://doi.org/10.5281/zenodo.10520670.

## Introduction

Single-cell RNA-sequencing (scRNA-seq) has become a popular approach for gaining valuable insights into various biological questions [Jovic et al., 2022, Sun et al., 2021, Hwang et al., 2018] at the cellular level. In most short-read scRNA-seq assays, such as the 10x Genomics Chromium 3′ system, cDNA reverse transcription is primed using oligo(dT) in order to capture the poly(A) tail of polyadenylated RNAs [10x, 2022a, Hrdlickova et al., 2016]. Then, the synthesized cDNAs are amplified, fragmented, and sequenced to generate sequencing reads as the readout of the RNAs primed by oligo(dT) primers. Because, in droplet-based scRNA-seq protocols, cells (or nuclei) are lysed within each droplet, all exposed adenine-single nucleotide repeats (A-SNRs) in lysed cells have the chance to be captured by oligo(dT) primers and generate “valid” reads. This is true regardless of whether or not the A-SNRs appear in a poly(A) tail, or internally within a spliced or unspliced molecule, and, of course, whether or not the underlying molecule is associated with a protein-coding or non-coding RNA. Oligo(dT) priming that occurs outside of the “expected” A-SNRs (i.e. the polyA tail of polyadenylated RNA molecules) is typically referred to as “off-target priming”. However, reverse transcription can be initiated by priming oligo(dT) at internal sites with as few as 6 consecutive As [Nam et al., 2002, Svoboda et al., 2022]. Additionally, a technical note from 10x Genomics [10x, 2021] describes and explains the mechanisms of generating sequencing reads from internal polyA sites on RNAs as well as on cDNAs, supporting the validity, and to some extent, prevalence, of off-target priming in scRNA-seq.

Recent studies have shown that off-target priming is prevalent in scRNA-seq. For example, a detailed technical note from 10x Genomics [10x, 2021] showed a high proportion of off-target reads across popular 10X scRNA-seq assays. Svoboda et al. [2022] showed the prevalence of intronic priming by analyzing publicly available datasets. He et al. [2023] processed eight 10X Chromium 3′ scRNA-seq datasets from both single-cell and single-nucleus samples to show that when considering all sense genic reads, up to 40% of unique molecule identifiers (UMIs) only have reads compatible with intronic regions in unspliced (or partially spliced) transcripts. In general, throughout this article, we will refer to “unspliced” molecules in the understanding that they may be actively undergoing splicing and hence be partially spliced. Meanwhile, researchers have realized the underappreciated value of off-target reads and designed sophisticated algorithms to incorporate off-target reads in scRNA-seq data analysis [Chamberlin et al., 2022, 10x, 2022b, Pool et al., 2023, Chari et al., 2023, Gorin et al., 2023]. One example is single-cell RNA velocity, in which cellular splicing dynamics are inferred using spliced gene counts from reads compatible with spliced transcripts, and unspliced counts accounting for reads compatible with unspliced transripts [La Manno et al., 2018]. Although these studies showed the existence of some types of off-target reads and suggested plausible biological interpretations, there is not yet a comprehensive study systematically analyzing all types of off-target reads from different genomic features and exploring their potential use cases.

In this study, we introduced `scCensus`, a comprehensive Nextflow [Di Tommaso et al., 2017] workflow for systematically classifying the off-target scRNA-seq reads from different genomic feature groups. We divided scRNA-seq reads into three categories: sense intragenic, antisense intragenic, and intergenic reads. We performed an in-depth analysis for reads belonging to each read group, and we observed that off-target reads from all genomic feature groups reflect meaningful biology. Our results show that intergenic scRNA-seq reads are enriched near open chromatin regions (OCR) as detected from single-cell sequencing assay for transposase-accessible chromatin (scATAC-seq), i.e., scATAC-seq peaks, and provide information about open chromatin regions. Furthermore, OCR-associated reads can result, at least at low resolutions, in clustering results consistent with the standard method. Furthermore, when sense and antisense intragenic reads are quantified separately we find that their quantification results are highly, but imperfectly, correlated, suggesting that antisense reads can be used to improve gene detection. On the other hand, some antisense reads are likely to be derived from genuine antisense transcripts [Pelechano and Steinmetz, 2013, Barann et al., 2013]. Finally, using splice-aware quantification methods [He and Patro, 2023, He et al., 2022], we find that the clustering results generated from spliced, unspliced, and ambiguous matrices are consistent to a large extent (at a coarse level), but also show informative differences (at finer granularity). Well-established marker genes of cell types were exclusively discovered from each of these count matrices, further suggesting that reads with different splicing statuses should be processed and analyzed separately and integrated in a later stage. All of these results suggest that off-target scRNA-seq reads reflect meaningful and interesting biology. Therefore, we urge the community to expand current analyses to incorporate such off-target fragments, and to develop novel methods and approaches that intrinsically account for such fragments.

## Methods

### Data Description

In this study, we processed seven scRNA-seq datasets generated using single cells and single nuclei samples from the brain, blood, and bone marrow of mouse and human. Details about the selected datasets can be found in Supp table 3. We processed all human datasets using the `GRCh38 version 2020-A` genome build and all mouse datasets with the `mm10 version 2020-A` genome build. Both genome builds were downloaded from the 10X Genomics website ^1^. For each genome build, we applied two sets of gene annotations, one was downloaded from the 10x Genomics website along with the genome build, and the other was the *scRNA-seq optimized gene annotations* [Pool et al., 2023]. The seven selected single-cell datasets span different species, tissue types, and sample sizes.

### Quantification

To perform downstream analyses, we processed the read alignment BAM files generated in Supplementary section A.2 for sense intragenic, antisense intragenic, and intergenic reads. The resulting count matrices represent the unique molecule identifiers (UMIs) that are compatible with sense intragenic (including intronic regions), antisense intragenic (including intronic regions), and intergenic regions. Intergenic regions are defined as genomic regions that do not intersect with any gene annotations. In the following text, we describe the quantification pipeline applied to each dataset.

First, we quantified the sense and antisense intragenic reads using `simpleaf` [He and Patro, 2023] to generate cell barcode-by-gene count matrices. For each single-cell dataset, we generated a total of six-count matrices for intragenic UMIs with a spliced transcript origin (*M*_S_), unspliced transcript origin (*M*_U_), ambiguous splicing status(*M*_A_), and UMIs from genes’ antisense, reverse complement strand with the three splicing statuses, 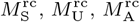, respectively. We note that only UMIs S U A that were not associated with any sense reads were used to generate the antisense count matrices (that is, sense assignment was preferred if it was possible), and the only change instructing `simpleaf` to generate antisense instead of sense count matrices was changing the `expected-ori` parameter from `fw` to`rc`.

The`simpleaf` pipeline used in this work involves two steps: reference index construction and sequencing read quantification. We used the *spliced+unspliced* (*spliceu*) reference [He et al., 2023], which contains the sequence of spliced transcripts and gene bodies. The gene body of each gene contains the contiguous genomic interval from the 5′ farthest exonic locus to the 3′ farthest exonic locus of each gene considering all its isoforms. Providing the *spliceu* reference to `simpleaf` triggered the USA mode [He et al., 2022] of the underlying `alevin-fry` module to generate three UMI count matrices, representing the UMI count of each gene in each cell with spliced, unspliced, and ambiguous splicing status. Briefly, when using *spliceu*, the spliced count matrix contains UMIs with exon-exon junctional mappings and without intronic alignments, the unspliced count matrix contains UMIs that are entirely or partially compatible with introns, and the ambiguous count matrix contains UMIs compatible with both spliced and unspliced transcripts, i.e., exonic UMIs.

Additionally, We developed a custom pipeline to process intergenic reads to generate four open chromatin region(OCR)- associated count matrices. In this work, we used the ATAC-seq peaks discovered from the ATAC-seq component of the single-cell multiome ATAC+RNA (scMultiome) assays to represent the experimental open chromatin regions. To show that our conclusions apply to unpaired ATAC-seq and RNA-seq data, we also used the peaks discovered from independent ATAC-seq samples for validation. In order to verify that the ATAC-seq peaks represent *cis*-regulatory elements, we apply the same pipeline using the candidate *cis*-regulatory elements (cCREs) from the SCREEN project [Consortium. et al., 2020] as the OCR features to generate cCRE-associated count matrices. As the same pipeline was applied to all OCR feature sets, in the following text we describe the pipeline for one feature set, *F*, which can be an ATAC-seq peak set or the cCREs from SCREEN. Note that in this pipeline, only UMIs that are not associated with intragenic reads were included to generate the count matrices.

The pipeline is divided into the following steps. First, we filtered the original feature set, *F*, into two subsets using `bedtools intersect` .The first subset, *F*_ig_, contains only intergenic features. The second subset, *F*_nc_, contains all features that do not intersect with protein-coding genes (on either strand of the genome). Second, we process all or a subset of sequencing data with *F, F*_ig_, or *F*_nc_ using the `bedtools closest` program by specifying `-d` and `-t first` to find the closest features for each read. Specifically, We used only intergenic reads together with *F*_ig_ to generate the intergenic feature-associated count matrix, 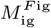. The subscript represents the reads (in this case, intergenic reads) used to generate the count matrix, and the superscript represents the feature set (in this case, intergenic features). nc

We used reads from UMIs that are not compatible with any protein-coding genes on either strand of the genome with *F*_nc_ to generate the non-coding features-associated count matrix, 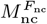. We used reads that are not compatible with protein-coding genes in the forward orientation with the whole feature set, *F*, to build the not-sense-coding feature-associated count nsc matrix, 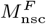. We used all read alignments with *F* to build an all all feature-associated count matrix 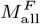. During this process, only alignments that are less than 4, 000 bases away from their nearest feature were used, corresponding to the reported length range of eRNAs and other lncRNAs [Wan et al., 2022].

We tested that using more restrictive thresholds for the distance did not affect our conclusions (data not shown). Finally, we processed all sets of filtered alignments in `Python` using `pysam2` to generate the feature-associated count matrices. We assigned each UMI to its closest feature, among all filtered alignments, to get the UMI count of each feature.

### Cell type identification

For each dataset, we used `sctype` [Ianevski et al., 2022] to assign a cell type to each high-confidence cell. The quantification results generated by `simpleaf` using the sense intragenic alignments were loaded into an `R` (version 4.3.2) environment as a `seurat` object [Hao et al., 2021] using the `loadFry` function from the `fishpond` Bioconductor package [Zhu et al., 2019]. For cell samples, we used the spliced and ambiguous total count to create the `seurat` object. For nucleus samples, we used the spliced, unspliced, and ambiguous total count to create the `seurat` object. As this is the standard strategy to create the count matrix for cell and nucleus samples, we call them the standard matrices *M*_std_ throughout the paper. For each `seurat` object, `SCTransform` [Hafemeister and Satija, 2019] with the default setting was used for preprocessing.

Next, we computed the top 100 principal components (PCs) by applying the `RunPCA` function using the variable features found by `SCTransform`. The number of significant PCs was found using `findPC` [Zhuang et al., 2022]. We then used these significant PCs to assign a cell cluster to each cell by calling `FindNeighbors` followed by `FindClusters`. The `resolution` parameter of `FindClusters` was set as 0.7. We then adapted the example code from the `sctype GitHub` repository^3^ to assign a cell type to each cell. The predicted labels were written to disk as a CSV file for future use. Although using a high clustering resolution, 0.7, might result in more cell clusters than the actual number of cell types, clusters with similar expression profiles should be assigned the same cell type by `sctype` [Ianevski et al., 2022].

### The sense intragenic read analysis pipeline

In this section, we describe the pipeline for analyzing the quantification results generated from the sense transcriptomic reads in each dataset using `simpleaf`. This analysis focused on comparing the cell clusters and the differentially expressed genes discovered from each cell type across different count matrices. The quantification results were loaded into R using `loadFry`.The spliced, unspliced, and ambiguous count matrices were saved separately.

We first created a `seurat` object for each of the following count types: (1) spliced counts (*M*_S_), (2) unspliced counts (*M*_U_), (3) ambiguous counts (*M*_A_), (4) spliced and ambiguous total counts (*M*_SA_), and (5) spliced, unspliced and ambiguous total counts (*M*_USA_). Usually, (4) and (5) are the standard count matrix for cell and nucleus samples. Each`seurat` object was processed as described in section 2.3. For each `seurat` object, we tuned the `resolution` parameter of `FindClusters` to find three cluster sets, each with a different number of clusters. Assuming that the number of cell types found by `sctype` (section 2.3) is *n*_*c*_, the three cluster sets have max(3, *n*_*c*_ *×* 0.2), max(7, *n*_*c*_ *×* 0.6), and *n*_*c*_, respectively. We performed this resolution sweep, as we want to see how the clustering changes and relates across different types of counts as the resolution of the clusters changes, as similarities present at a coarse resolution may either persist or diminish at a finer resolution. To assess the similarity of two sets of cluster assignments with the same number of clusters, we adopted the evaluation metrics used in Yu et al. [2022], namely the adjusted Rand index (ARI) [Chiquet et al., 2023], normalized mutual information (NMI) [Chiquet et al., 2023], and Fowlkes–Mallows index (FMI) [Galili, 2015]. All three metrics range from 0 to 1, where 1 represents a perfect match.

We then computed the differentially expressed genes (DEGs) for each cell cluster identified by `sctype` (section 2.3) in each `seurat` object using the `FindAllMarkers` function from `seurat` with the default setting.

### The antisense intragenic read analysis pipeline

In this analysis, we explored the count matrix containing antisense intragenic UMIs (*M*_anti_). The antisense count matrix was generated by quantifying UMIs that do not have any associated sense intragenic alignments across all splicing statuses (section 2.2) and loaded into `R` using `loadFry`. We also generated a sense count matrix by considering the sense UMIs across all splicing statuses, same as the *M*_USA_ described in section 2.4. Then, based on *M*_sense_ and *M*_anti_, we *imputed* genes that were likely to be missing from the *M*_sense_ matrix to generate an imputed sense count matrix, *M*_imputed_. This was done by identifying cell, gene pairs whose count in *M*_sense_ was zero, but whose count in *M*_anti_ was non-zero; these cell, gene pairs were then assigned a value of 1 in *M*_imputed_. All non-zero cell, gene entries from *M*_sense_ were carried over directly to *M*_imputed_ without modification. A sense and antisense total count matrix, *M*_genic_, was also created by summing *M*_sense_ and *M*anti.

Next, we computed the correlation of each cell in *M*_sense_ and *M*_anti_ using all genes that are detected in that cell in the *M*_sense_ matrix. We also tried to use genes that are detected in *M*_anti_ of that cell, and genes detected in both *M*_sense_ and *M*_anti_. The correlation was assessed using Spearman’s rank correlation coefficient (*ρ*), the Pearson correlation coefficient (*r*), and the cosine similarity (cos).

Then, we applied the same clustering analysis and differential expression analysis pipelines introduced in section 2.4 to *M*_sense_, *M*_anti_, *M*_imputed_, and *M*_genic_, to find the cell clusters and the DEGs discovered in the `sctype` cell types (section 2.3).

### The intergenic read analysis pipeline

In this section, we describe the analysis pipeline used for analyzing intergenic scRNA-seq reads (section 2.2) from each dataset. The input of this analysis consists of an intergenic count matrix *M*_int_, an intergenic open chromatin region-associated count matrix *M*_ocr_, as well as a standard count matrix, *M*_std_, generated from sense intragenic reads (section 2.3). The *M*_std_ of cell samples were generated by summing the spliced and ambiguous count matrices, and the *M*_std_ of nucleus samples were generated by summing the spliced, unspliced, and ambiguous counts. We explored all types of OCRs defined in 2.2, including the intergenic OCRs, not-protein-coding OCRs, non-sense-coding OCRs, and all OCRs from ATAC-seq peaks and SCREEN cCREs. Those count matrices were loaded into R using either `loadFry` or the `Read10X` function from `seurat`, as appropriate. We built a `seurat` object for each count matrix separately and used them in the following steps.

To evaluate the abundance of intergenic reads near ATAC-seq peaks, we designed a statistical test to evaluate if the ratio of the count of intergenic regions from *M*_int_ to its OCRs from *M*_ocr_ is significantly different than the ratio of their size (section A.3).

The intergenic read analysis pipeline contains two components. First, we performed a clustering analysis as discussed in section 2.4 using *M*_std_, *M*_ocr_, and *M*_int_. For *M*_std_, we set the number of variable features as 3, 000 when invoking `SCTransform`. For *M*_ocr_, we used the top 70% of features ranked by their variance as the variable features, because of the sparsity. Meanwhile, we applied the standard `Seruat` pipeline (using `NormalizeData, ScaleData`, and `FindVariableFeatures` sequentially) instead of `SCTransform` for all count matrices except *M*_std_, because (i) running `SCTransform` with too many variable features was computationally prohibitive, (ii) we processed *M*_std_ using `SCTransform` in all other sections, and (iii) the main purpose of this step is to test if the clusters generated from *M*_std_ are similar with others, which should not happen owing to processing data differently.

Second, we showed that, in the multiomics datasets (Supp table 3), the ATAC-seq peak counts of a cell are more similar to its RNA-seq OCR counts than the RNA-seq OCR counts of most other cells. In this analysis, because the processed datasets are single-cell multiome ATAC + Gene Expression (scMultiome) datasets, the two modalities of each dataset — represented by the RNA-seq OCR count matrix and the ATAC-seq peak count matrix — contain the same set of features (ATAC-seq peaks) but different types of readouts. For each cell, its RNA-seq OCR counts contain the UMIs representing the RNAs that have read alignments associating with ATAC-seq peaks, while its ATAC-seq peak counts are from the readout of the DNA fragments measured by ATAC-seq. This allows us to ascertain ground truth correspondence, against which we can then compare by evaluating the similarity of the RNA-seq OCR and ATAC-seq peak counts. The ATAC-seq peak count matrix was downloaded as a part of each scMultiome dataset and loaded into R using `Read10X`. It consists of the same set of features (peaks) and cells with the OCR count matrix.

Cosine similarity was used to evaluate the association of the two modalities to avoid the effect of genes that are undetected in both modalities. The null hypothesis is that the ranking of the cosine similarity of each cell’s counts across the two modalities should not differ from the ranking of the cell’s ATAC-seq peak count to a random cell’s OCR count. In other words, there should be no meaningful association between the ATAC-seq peak counts in a cell and the RNA-seq OCR peaks in the same cell, so that, if we ranked one according to the other with respect to their cosine similarity, it would appear at an essentially random position in the ranked list. The statistical significance of the rankings was evaluated by a Wilcoxon rank sum test, the lower-tailed t-test, and the Kolmogorov–Smirnov test on the two distributions.

## Results

Below, we describe the results of analyzing seven selected single-cell datasets (Supp table 3). First, we examine the selected single-cell multiome ATAC+Gene expression (scMultiomics) datasets and show that intergenic RNA-seq reads retain cellular chromatin structure, and can be used for identifying similar cells across modalities. Next, we explain the relationship between the count matrices generated from sense and antisense intragenic reads, and the possibility of using antisense reads to augment the standard count matrix and to study antisense transcription. Finally, we discuss the clustering and differential expression analysis results using sense intragenic reads from different splicing statuses.

The first striking observation from our analyses is that off-target priming is prevalent in scRNA-seq data. As shown in Supp table 3, among the seven scRNA-seq datasets we analyzed— spanning different species, sample types, tissue types, and sample sizes — on average 78% of sense intragenic UMIs are not exclusively compatible with spliced transcripts (i.e. do not span an exon-exon junction), and this percentage can grow up to up to 96% in some datasets. Antisense transcript UMIs on average account for 18% (maximum 30%) of intragenic UMIs. Intergenic UMIs on average account for 8% (13% in maximum) of total genomic UMIs. This work aims to explore the biological interpretability of off-target scRNA-seq reads and shed light on the potential usages of off-target reads to improve and expand current single-cell analysis methods.

### Intergenic reads associate with cell-specific open chromatin regions

Intergenic reads are reads compatible with the genome but not gene annotations. In the processed datasets, intergenic reads on average account for approximately 10% of the high-quality reads (Supp table 3). Although those reads do not match any existing gene annotations, they are presumed to arise from RNAs, and to represent cellular transcripts. In `scCensus`, we associate scRNA-seq reads with *intergenic* open chromatin regions (OCRs) detected from single-cell sequencing assays for transposase-accessible chromatin (scATAC-seq), i.e., scATAC-seq peaks.

In this section, we present our intergenic read analysis results using the selected scMultiome datasets, in which the gene expression and chromatin accessibility profiles of each cell are measured by scRNA-seq and scATAC-seq, respectively. Briefly, we counted the intergenic scRNA-seq UMIs associated with each intergenic ATAC-seq peak, to generate a scRNA-seq OCR count matrix *M*_ocr_. In this count matrix, the features represent the intergenic ATAC-seq peaks, and the entries represent the scRNA-seq UMI count of each feature. Similarly, we generated an intergenic count matrix, *M*_int_, by counting the UMIs compatible with the intergenic regions between each pair of adjacent genes. We compared the results from *M*_ocr_ and *M*_int_ with the standard count matrix *M*_std_ (section 2.6). We focus here on the results from a human PBMC scMultiomics dataset, the results from other datasets and other types of OCR count matrices are listed in Supp figs. 5 to 15.

We first validated the biological interpretability of intergenic reads to show that they *do not* appear to be DNA contamination, ambient RNAs, or ribosomal RNAs. Our results demonstrate that for each cell, the size ratio of intergenic regions to intergenic OCR regions was on average six times higher than their count ratio, with a *p*-value *<* 10^−8^(Supplementary Files). The *p*-value was computed using a one-side t-test (section 2.6). Moreover, the cell clusters obtained from *M*_int_ and *M*_ocr_ were consistent with those obtained from the standard count matrix *M*_std_, especially when the clustering resolution was low, as shown in figs. 1a, 1b and 24 to 30). This suggests that, at the coarsest resolutions, the large-scale similarity structure of the cell count matrices persists, even when vastly different types of features are being quantified.

**Fig. 1.**
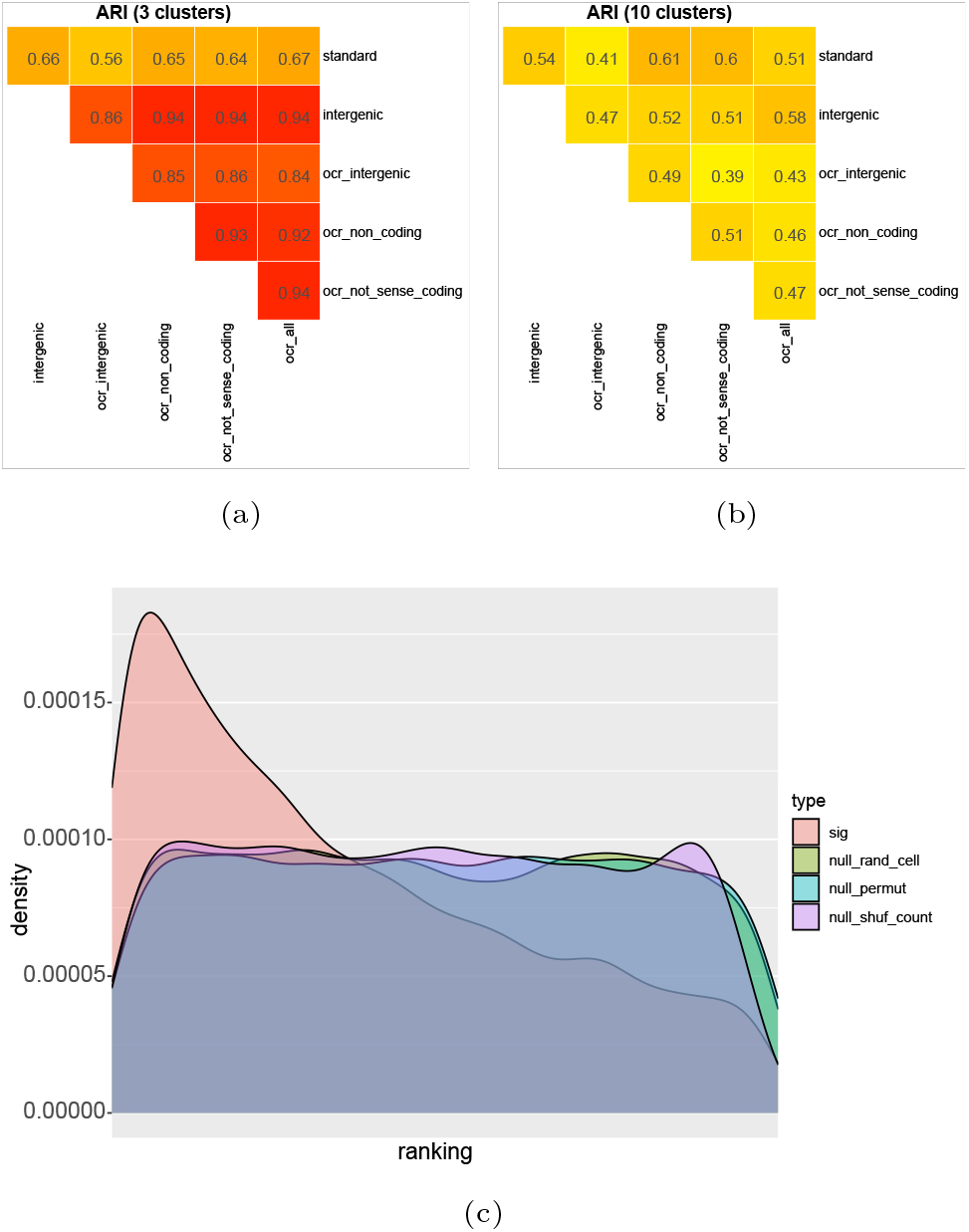
Intergenic reads reveal cellular chromatin structure. ARI stands for Adjusted Rand Index. ARI=1 means the compared sets of cluster assignments are identical. *ocr intergenic* means results from the intergenic OCR count matrix. *intergenic* means the intergenic count matrix. *ocr non coding* means the not protein-coding OCR count matrix. *ocr not sense coding* means the OCR count matrix containing all but not reads from the sense orientation of protein-coding genes. *ocr all* means the total OCR count matrix. Panels (a) and (b) show the ARI of the clustering results using low and high resolution, respectively. Section 3.1 discussed the ARIs of *ocr intergenic* and *intergenic*. Panel (c) shows the ranking of the cosine similarity of *ocr intergenic* to ATAC-seq counts of the same cell compared with all other cells (*sig*), together with three null distributions to show the significance of the high ranking (section 3.1).

The similarity was measured using the Adjusted Rand Index (ARI) [Chiquet et al., 2023]. An ARI = 1 means that two sets of cluster assignments are identical. When the clustering resolution is low, the cluster assignments under the *M*_int_ and *M*_ocr_ matrices are very similar to the standard count matrix, with an ARI higher than 0.9. As the clustering resolution increases, the ARI decreases. Although it is uncertain whether the divergence in the high resolution was caused by the high sparsity of *M*_ocr_, the high similarity at a low resolution provides compelling evidence that *M*_ocr_ contains sufficient biological signals to distinguish the major cell types. Similar results were obtained when performing our analysis using independent scATAC-seq datasets, or using a reference that augments unannotated 3′ UTRs [Pool et al., 2023] (Supplementary Files and Supp table 5). These two pieces of evidence together show that intergenic reads probably reflect meaningful biological signals of the underlying cells, and that the signal-to-noise ratio is sufficient to extract some of this information. One likely explanation for these fragments is that they originate from non-coding RNAs originating from intergenic open chromatin regions (e.g. enhancer RNAs [Sartorelli and Lauberth, 2020] and promoter RNAs [Chellini et al., 2020]).

Furthermore, by comparing *M*_ocr_ with the standard scATAC-seq count matrix, we discovered that intergenic reads reveal cell-specific chromatin structure. Briefly, for each cell *i*, we computed the cosine similarity of its intergenic ATAC-atac seq peak counts 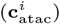to the intergenic OCR counts of every cell *j* 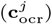. We define these similarities as sim cos 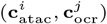, where *i, j* ∈ {0..*N −* 1} and *N* is the number of cells in the dataset. Our results showed that if we rank the cosine atac similarity of the same cell across modalities —cos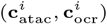 — the distribution is heavily skewed to the left (toward low ranks). That is, on average, the cosine similarity of the same cell across modalities is much higher than the similarity of cell *i* with most other cells *j*. This distribution is shown as the curve labeled as *sig* in Figure 1c. We considered several different null models. Null distributions were generated by atac evaluating cos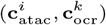 for a random cell *k* for each *i* (with replacement), shuffling the two count matrices, and permuting the ranking of cells, corresponding to the curves labeled as *null rand cell, null shuf count*, and *null permut* in fig. 1c. The difference of *sig* compared with the three null distributions is statistically significant, as evaluated by a Wilcoxon rank sum test, with *p*-values all *<* 10^*−*16^. Although we focused on the intergenic reads and peaks in this analysis, the same conclusions still held when including more ATAC-seq peaks, as shown in Supp figs. 17 to 23 and Supplementary Files.

In conclusion, by counting the intergenic scRNA-seq UMIs proximate to intergenic scATAC-seq peaks, we find that intergenic scRNA-seq reads are enriched around scATAC-seq peaks and can produce cluster assignments consistent with those from standard scRNA-seq count matrix, indicating their biological interpretability. Perhaps even more interesting, the strong association between intergenic scRNA-seq and scATAC-seq peak counts suggests that intergenic scRNA-seq reads reflect cell-specific chromatin structure, indicating that they might originate from intergenic regulatory RNAs, such as enhancer RNAs [Sartorelli and Lauberth, 2020, Young et al., 2017] and promoter RNAs [Chellini et al., 2020]. The strong association also suggests the possibility of designing novel distance metrics according to these two modalities to help in aligning or integrating unpaired scATAC-seq and scRNA-seq.

### Antisense transcriptomic reads contain mixed signals from mRNAs and regulatory RNAs

In the context of scRNA-seq, antisense intragenic reads are reads that map to gene annotations in an antisense manner. These reads are usually discarded in assays such as 10X Chromium V2 and V3 where stranded sequencing protocols are used and the reads are expected to align with the underlying genes in the forward (or sense) orientation. In our processed datasets, antisense intragenic UMIs can account for up to 30% of total intragenic UMIs (Supp table 3), with an average of 18% across the datasets we evaluated. 10x Genomics previously demonstrated the prevalence of antisense scRNA-seq reads [10x, 2021], and also discussed four potential mechanisms that can explain their presence in the collection of sequenced molecules, including priming by the template-switching oligo, poly(dT) primer strand invasion, first-strand cDNA priming, and sense-antisense fusion. Antisense reads may also arise from RNAs generated via antisense transcription [Katayama et al., 2005, Barman et al., 2019, Barann et al., 2013] or other types of bidirectional transcription events [Morris et al., 2008].

In this section, we discuss our findings by comparing the analysis results obtained from the sense (*M*_sense_) and antisense (*M*_anti_) intragenic count matrices of each dataset (Supp table 3). The antisense count matrix contains intragenic UMIs only associated with antisense fragments (section 2.5). We focus here on the results from a human brain scMultiomics dataset, the results from other datasets are listed in Supp figs. 17 to 33.

We first compared the clustering results obtained from the sense and antisense counts. As shown in fig. 2a, we observe an ARI of 0.65 between the cluster assignments from sense and antisense count matrices when both have 11 clusters — the number of cell types discovered by `sctype` (section 2.3). This indicates that antisense reads contain interpretable and meaningful biological signals (by virtue of their substantial overlap with the intended signal from the sense intragenic reads). Then, we confirmed that some antisense reads are the likely products of the four technical mechanisms discussed above. Because the four technical mechanisms can happen only *after* valid RNA priming, the resulting antisense reads should also reflect gene expression, and, therefore, their quantification results should be well-correlated with their sense counterpart. That is, while we expect antisense counts to diverge from sense counts, they nevertheless require the actual presence, in the assayed cell, of the RNA molecule to which they map. Our results show a moderate-to-high correlation between cells’ sense and antisense counts. Among the three selected metrics, the Spearman *ρ* (fig. 2b) correlation coefficient of cells’ sense and antisense counts are centered around 0.4, the Pearson *r* (fig. 2c) centered at 0.6, and the cosine similarity(fig. 2d) are centered around 0.7. We also confirmed that the correlation scores are very close to zero when using shuffled sense and antisense count matrices (Supp fig. 16).

**Fig. 2.**
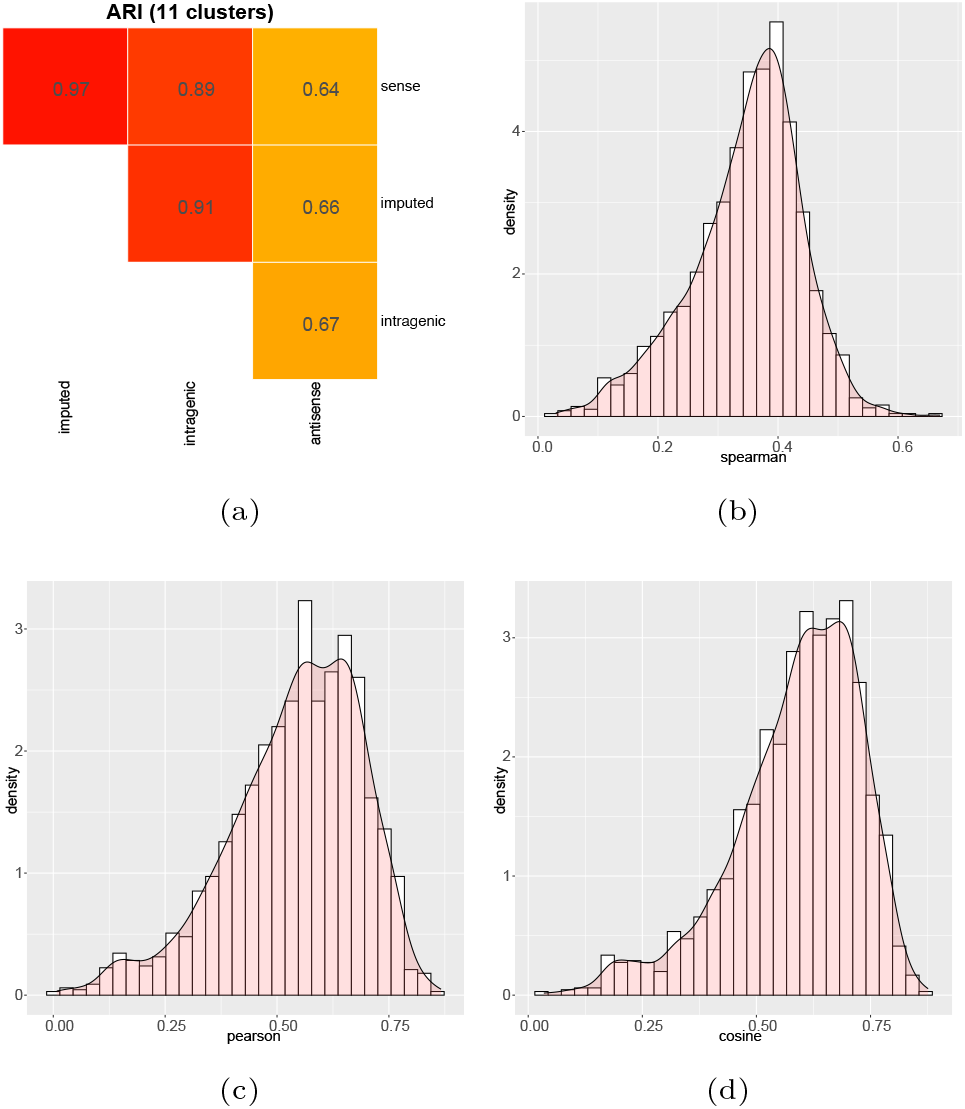
Antisense intragenic reads are interpretable and show a moderate-to-high correlation with sense reads. Panel (a) shows the high Adjusted Rand Index scores of the cluster assignments obtained from *sense, imputed, antisense*, and *intergenic* count matrix (section 3.2). ARI=1 means the compared sets of cluster assignments are identical. Panels (b), (c), and (d) show the Spearman *ρ*, Pearson *r*, and cosine similarity of the sense and antisense counts of cells, respectively.

Our results suggest that, on one hand, many antisense reads seem to directly provide evidence for the presence of RNA molecules in cells that agree with the counts obtained in the sense matrix. Thus, this substantial fraction of the antisense reads is compatible with the four technical mechanisms previously described in the 10x Genomics technical note. On the other hand, the imperfect correlation also suggests that some fraction of antisense reads appear not to be explained by these specific technical mechanisms. Together with the imperfect ARI discussed above, as well as the distinct cell markers discovered from the sense and antisense count matrices (Supp figs. 31 to 33), we hypothesize that some antisense reads likely arise from antisense transcription or other bidirectional transcriptional activities.

In addition to evaluating the concordance between the sense and antisense intergenic count matrices, we also explored the possibility of using the information contained in the antisense count matrix to augment (i.e. improve the sensitivity and reduce the sparsity [Bouland et al., 2023] of) the standard (sense) scRNA-seq count matrix.

Due, in part, to limited sampling from a finite population of molecules (and potentially other sources of increased “dropout” in scRNA-seq [Qiu, 2020]), the standard count matrix is dominated by zeros, i.e., it is a very sparse matrix. As we observed the correlation between cells’ sense and antisense counts, but note that antisense reads are usually excluded from the standard count matrix, we attempted two relatively naïve methods to improve the sensitivity of *M*_sense_ according to *M*_anti_.

The first strategy examines all entries that have a non-zero entry in *M*_anti_, and if the corresponding entry in *M*_sense_ is zero, it is changed to a 1. This results in the generation of an augmented sense count matrix, *M*_imputed_ (section 2.5). We only change zeros to ones instead of copying over the actual antisense counts to reduce the potential disturbance caused by the signals from regulatory RNAs in *M*_anti_.

The second strategy sums *M*_sense_ and *M*_anti_ matrices to obtain an intragenic count matrix, *M*_genic_, similar to considering reads in both orientations when generating the standard count matrix. Both strategies led to a substantial reduction of sparsity. Specifically, if one considers matrix entries where *M*_sense_, *M*_anti_, or both are non-zero, then in 19.3% of such cases we observe a corresponding 0 entry in *M*_sense_ and a non-zero entry in *M*_anti_ across the processed datasets. In other words, among the non-zero entries of *M*_genic_, 19.3% are zero in *M*_sense_ but non-zero in *M*_anti_. We note that our strategies primarily aided in the detection of the presence of genes in cells where they were not previously detected, but not in the detection of genes not detected across any of the cells in *M*_sense_ (only less than 0.2% of the values imputed from *M*_anti_ were for genes not otherwise present when looking across all cells in *M*_sense_). Again, this behavior supports the sense and antisense reads reflecting mostly the same underlying biology. We found that the augmenting strategies reduced the count sparsity in the scRNA-seq matrices, but did not substantially “disturb” or alter the results of standard analyses. Figure 2a showed that both strategies yielded cluster assignments consistent with the sense count matrix. The ARI of the results from *M*_sense_ to *M*_imputed_ is 0.97, somewhat higher than the ARI from *M*_sense_ to *M*_genic_ of 0.89.

In conclusion, by quantifying sense and antisense intragenic scRNA-seq reads separately, and comparing their analysis results, we found a moderate correlation between their counts and a moderate-to-high similarity between their cluster assignments. Our results reinforced that a substantial fraction of the antisense reads observed in scRNA-seq are the result of specific technical artifacts that nonetheless reflect the expression of genes that may otherwise be detected in the sense orientation by a more sensitive assay or deeper sequencing. We showed that incorporating the counts generated by antisense reads can help reduce the sparsity and improve the sensitivity of gene detection. Moreover, another fraction of antisense reads seem to likely originate from regulatory RNAs derived from antisense transcription [Katayama et al., 2005, Barman et al., 2019, Barann et al., 2013] or other types of bidirectional transcription events [Morris et al., 2008]. Proper incorporation and assessment of these reads deserves its own dedicated analysis, and likely the development of novel methods to account for them in preprocessing and processing.

### Sense intragenic reads with different splicing statuses reveal different transcriptional information

Intronic reads are the reads compatible with annotated gene models but only with their intronic regions, i.e., unspliced transcripts.

One mechanism for generating such off-target reads is intronic polyA priming [10x, 2021]. Such priming occurs as many introns contain short (and even moderate-length) polyA tracts, and evidence has shown that polyA tracts of length 6 to 8 are sufficient to be anchored by oligo(dT) primers [Nam et al., 2002, Svoboda et al., 2022]. In our selected datasets, less than 40% sense intragenic UMIs can be unambiguously classified as arising from spliced transcripts (Supp table 3). We note, however, that we have adopted in this manuscript a rather conservative notion of spliced and unspliced status [Eldj’sarn Hjörleifsson et al., 2022, He et al., 2023], and other approaches, that consider purely exonic reads as arising from spliced RNA, will lead to different ratios. Existing studies have shown the prevalence of intronic reads [He et al., 2023, Pool et al., 2023, Chamberlin et al., 2022, 10x, 2021] and proposed algorithms that utilize unspliced reads in different ways [La Manno et al., 2018, 10x, 2022b, Gorin et al., 2023].

In this section, we focus on explaining some caveats with the currently common approaches for utilizing intronic reads for standard types of analysis, as well as interesting findings that underscore the necessity of expanding existing, and designing novel, algorithms to consider signals from both splicing statuses separately. Specifically, sense intragenic reads are quantified into three count matrices, *M*_S_ contains UMIs only compatible with spliced transcripts, *M*_U_ contains UMIs compatible only with unspliced transcripts, and *M*_A_ contains UMIs compatible with both, i.e., having an ambiguous splicing status. Our analysis also included two combinations of the three-count matrices. *M*_SA_ was created by summing *M*_S_ and *M*_A_ matrices elementwise, and *M*_USA_ was created by summing *M*_U_, *M*_S_, and *M*_A_ matrices elementwise. Usually, *M*_SA_ is used as the standard count matrix for datasets from single-cell samples, and *M*_USA_ is used for single-nucleus samples. We focus on the results from a human PBMC scMultiome dataset, the results from other datasets are listed in Supp figs. 34 to 47

Our clustering analysis results showed that, when using a coarse resolution clustering to assign cells into major cell types, like B cells, T cells, and Monocytes in PBMC [Sen et al., 2018], the evaluated count matrices all resulted in consistent cluster assignments, as demonstrated by the very high ARIs displayed in fig. 3a. This suggests that the difference between major cell types in these matrices is robust, so that they can be easily distinguished from all tested combinations of splicing statuses. The cluster assignments from *M*_USA_ had slightly lower ARIs than others; this may be caused by the specific distance metric or the random seed used for finding cell clusters [Hao et al., 2021], as the clusters under this count matrix became more concordant when using a slightly higher resolution (Supp fig. 35b). The increment of ARIs as the clustering resolution increases for *M*_USA_ when other ARIs are all decreased also highlights the need to take more than one of the count matrices as input to validate the final clustering results.

**Fig. 3.**
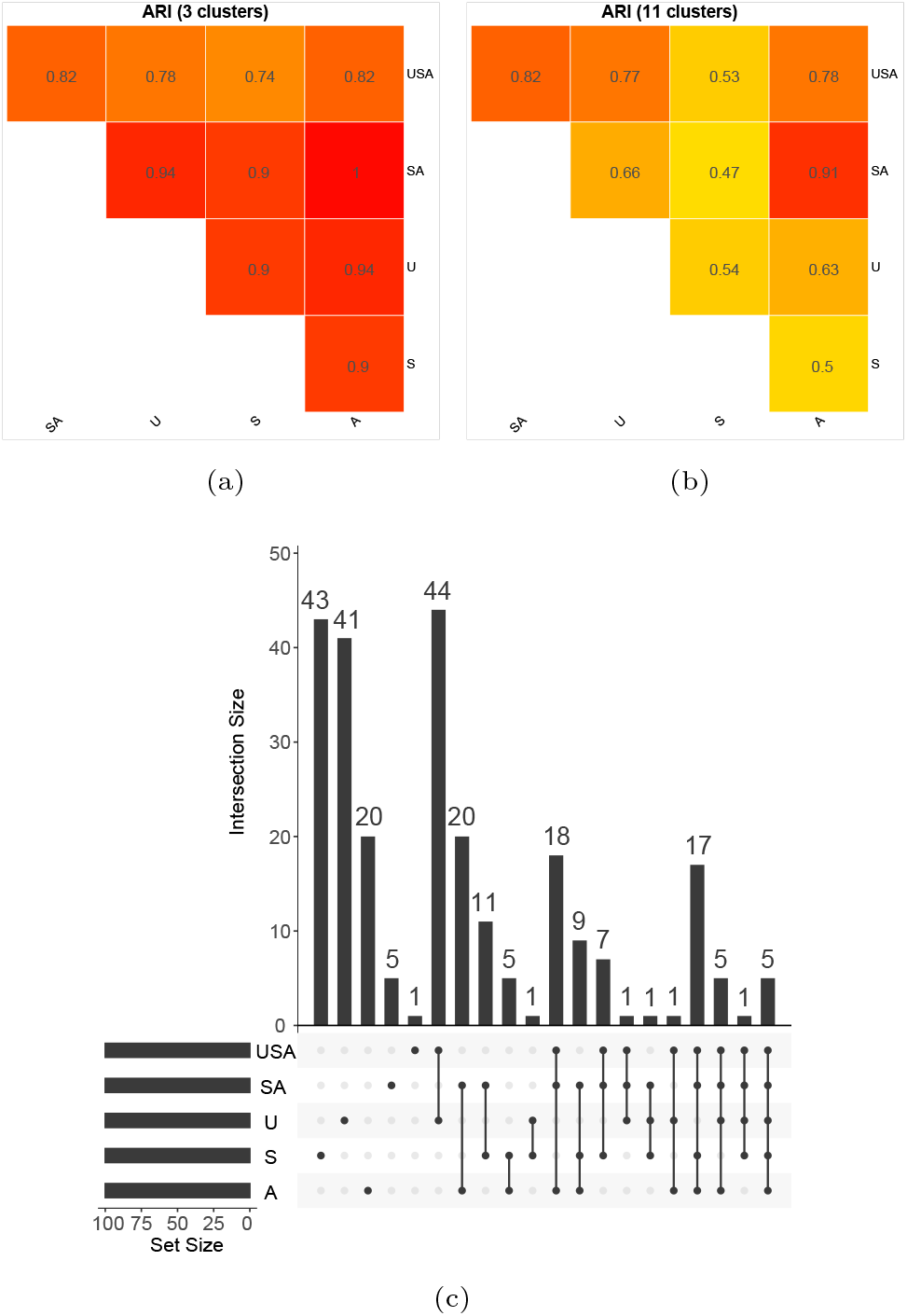
Sense intragenic reads with different splicing statuses contain distinct signals. *U* denotes the analysis results from the unspliced counts, *S* denotes results from the spliced counts and *A* denotes results from the ambiguous counts. *USA* is the sum of *U, S*, and *A* and *SA* is the sum of *S* and *A* (section 3.3). Panels (a) and (b) show the high Adjusted Rand Index scores of the clustering results generated from the count matrices using a low and high clustering resolution, respectively. ARI=1 means the compared sets of cluster assignments are identical. Panel (c) shows the intersection of the DEGs discovered from hippocampus neuron cells from the count matrices.

Furthermore, our clustering analysis results showed that, when using a high resolution (i.e. a fine-grained clustering) as in the standard clustering analysis pipeline [Hao et al., 2021], the cluster assignments from different combinations of splicing status show basic consistency but also express interesting differences, with ARIs ranging from 0.5 to 0.85. These imperfect ARIs suggest that, when using a single mixture of counts to perform clustering analysis, no matter which combination is used, the pipeline tends to cluster cells based on the outstanding signals from that combination, and interesting, potentially divergent, signals from other combinations remain latent.

The difference of the cluster assignments from *M*_S_ and *M*_U_ is especially important when clustering cells under active differentiation. For example, although cells under active differentiation should show consistent gene expression profiles under the spliced counts, cells in an early stage might have a distinct gene expression profile under the unspliced counts compared with cells in the terminal stage, because the spliced and unspliced transcripts in cells are desynchronized because of the kinetics of splicing [Alpert et al., 2016]. The difference of the cluster assignments of *M*_SA_ and *M*_USA_ in fig. 3b emphasizes the inconsistency by simply using *M*_USA_ as the standard count matrix for datasets from both single-cell sand single-nucleus samples, as e.g., `CellRanger` [Zheng et al., 2017] does in versions *≥* 7.

Our differential expression analysis results suggested that in each cell type, the counts from the three splicing statuses resulted in largely non-overlapping differentially expressed genes (DEGs), as shown in fig. 3c. For spliced and unspliced counts, over 40 of their top 100 DEGs are exclusive. As for the two combinations, *M*_USA_ and *M*_SA_, their top 100 DEGs are the mix of the DEGs obtained from *M*_U_, *M*_S_, and *M*_A_ separately. A closer look at these DEGs reveals that, as we concluded from the clustering analysis results, the DEGs obtained from each combination of splicing status contain exclusive, well-documented, cell type markers (Supplementary Files). For example, *CST3*, an important marker gene for monocytes [Hu et al., 2020], was identified as a DEG for the classical monocyte cell type when using *M*_S_ and *M*_A_, but does not show up among the top DEGs when using the two combinations, *M*_SA_ and *M*_USA_. Similarly, *CD86*, a marker gene for Dendritic cells [Tze et al., 2011], was found as a DEG of the Myeloid Dendritic cell type when using *M*_U_, but in none of the other modalities. One more example of the marker genes that were only discovered from *M*_S_ is a well-known *CD8* + T cell marker, *CD3G* [Li et al., 2019]. All these exclusive marker genes discovered using the counts of the three splicing statuses separately, suggest that simply summing them together to generate the standard count matrix might not be the most appropriate way to utilize the information from different splicing statuses.

In conclusion, by quantifying sense intragenic scRNA-seq reads according to their splicing status and performing clustering analysis and differential expression analysis on them and their combinations, we found that their cluster assignments slightly diverge but still show consistency and well-known cell type markers are discovered exclusively from each modality. Together with the previous finding that unspliced counts show gene length bias [Phipson et al., 2017, 10x, 2021, Chamberlin et al., 2022, Gorin et al., 2023], our results emphasize the necessity of improving existing methods and developing new algorithms to consider the signals from different splicing statuses jointly, but separately, for more comprehensive analysis results.

## Discussions

In this work, we have analyzed, across different organisms, annotations, and tissue types, off-target scRNA-seq reads compatible with intergenic, sense intragenic, and antisense intragenic regions. Our results draw a holistic picture of the off-target scRNA-seq reads, evaluate their biological interpretability, and show examples of using off-target reads to improve single-cell analysis from different perspectives.

Specifically, our intergenic read analysis results suggest that intergenic reads are likely to arise from regulatory RNAs, such as enhancer RNAs and promoter RNAs. We showed that intergenic reads reflect the chromatin architecture of cells, and have a strong association with scATAC-seq data, indicating the possibility of using them for aligning or integrating unpaired scRNA-seq and scATAC-seq data.

Furthermore, our antisense read analysis results indicate that antisense reads contain mixed signals relating to both gene expression and regulation. We show that antisense reads can be used to reduce the sparsity and increase the sensitivity in the scRNA-seq count matrix, and they also have the potential to provide insights into antisense transcription and other bidirectional transcription events.

Finally, we have analyzed sense intragenic reads from different splicing statuses separately and find that reads with different splicing statuses contain distinct signals signals, and, in part, signals from different stages of transcription. We found that the results of the clustering analysis and differential expression analysis using reads from different splicing statuses and their combinations are largely consistent but also reflect interesting disagreements. Especially interesting was that well-established cell markers can be found in each modality exclusively. Our result highlighted the necessity of expanding current methods and designing new analysis algorithms that can consider signals from different splicing statuses jointly, but individually, to draw more comprehensive biological conclusions.

Some constraints we adopted in our analysis highlight the limitations and future directions of this work. First, because of the limited sensitivity and relatively low number of UMIs per gene, and the limited read lengths (most reads are 100 bases long) in scRNA-seq protocols we evaluated, in this work we have assumed reads compatible with introns to arise from unspliced transcripts, and reads compatible with exon-exon junctions to arise from spliced transcripts, ignoring the specific category of partially-spliced transcripts. Yet, given the relevant kinetics and speed of splicing, it is likely that a substantial fraction of measured molecules are actually partially spliced. An alternative assay, such as long-read single-cell sequencing, may provide more insight into the underlying splicing dynamics, and may provide a greater ability to properly categorize read evidence arising from partially spliced molecules. Moreover, we assigned an ambiguous splicing status when unable to determine the splicing status of reads. One future direction is to develop more sophisticated methods to resolve the splicing status ambiguity, or to assign a meaningful probability to such a status [He et al., 2023].

Second, we defined intergenic and intronic regions according to existing gene annotations, but some scRNA-seq reads may arise from unannotated transcripts and thus may be assigned an incorrect genomic feature type. One future direction is to improve existing gene annotations, as discussed in Pool et al. [2023] and Barquin et al. [2023], to reduce such misclassification.

Most importantly, in this work, we focused on analyzing the impact and and potential uses of the latent signals encoded within off-target scRNA-seq reads using simple examples and strategies. This is possible, in part, because such signals look to be substantial and relatively strong. In the end, however, the proper way to incorporate and integrate such signals is to develop methods and algorithms that include and model them from the start, and propagate the relevant associated information through the entire single-cell and single-nucleus preprocessing and processing pipelines. From this perspective, the current work is not only an analysis of the prevalence and characteristics of off-target reads, but a motivation and call-to-arms of the single-cell method development community to design algorithms, analysis methods, and software tools to more comprehensively and holistically model and make use of these off-target reads.

## Footnotes

1 https://support.10xgenomics.com/single-cell-gene-expression/software/downloads/7.0/

2 https://github.com/pysam-developers/pysam

3 https://github.com/IanevskiAleksandr/sc-type

