## Supplementary material for "scCensus: Off-target scRNA-seq reads reveal meaningful biology": Supp

### Appendices

#### Appendix A Supplementary Methods

##### A.1 The transcript composition analysis pipeline

In this analysis, we explored the composition of transcripts in each type of gene annotation. The results of this analysis can be found in section B.1 and Supplementary files. We aim to explain that because of the prevalence of internal adenine-single nucleotide repeats(A-SNRs) in all types of RNAs, and the fact that exons are shared by spliced transcripts and their unspliced precursor, the splicing status of the RNA origin of sequencing reads and their UMIs are usually hard to determine. The analysis was carried out using an R version 4.3.2 environment. The analysis was carried out for each pair of genome and gene annotation sets (either the standard gene annotations or the *scRNA-seq optimized gene annotations*). In the following, we describe the analysis for one genome, gene annotation set pair.

Briefly, the gene annotation GTF file was loaded into R as a `TxDb` object using the `makeTxDbFromGFF` function from `GenomicRanges` (Lawrence et al., 2013). The genome FASTA file was loaded using the `readDNAStringSet` function from `Biostrings` (Pagès et al., 2023). The exon, transcript, and gene annotations were extracted from the `TxDb` object using `exonsBy`. `transcripts` and `genes` from `GenomicFeatures`, respectively. Each transcript's terminal kilobase was extracted by retracing the exonic bases from the $3'$ end until reaching the $1,000$-th exonic base or the $5'$ end. Each transcript's introns were defined as the set difference between the transcript body and its exons using the `setdiff` function from `GenomicRanges`.

Subsequently, the cardinality of exons and the size of the $3'$ terminal exon of each transcript were obtained by processing the transcript annotations. The cardinality of poly-A and poly-T tracts in exons, spliced transcripts, unspliced transcripts, and the $5'$ terminal kilobase of introns were examined using the `vmatchPattern` function from `Biostrings` to find the A-SNRs of the length 6 bases and more.

##### A.2 The read alignment analysis pipeline

In this analysis, we explored the cardinality of scRNA-seq reads compatible with the feature categories we are interested in. We performed this analysis for each of the seven selected single-cell datasets independently for each set of gene annotations (Section 2.1). The results of this analysis can be found in section B.2 and Supplementary files. In the following, we describe the procedure applied to each dataset.

The input data of this analysis consists of a genome-coordinate read alignment BAM file and a set of genomic annotation BED files, each containing the feature records from a feature category. To save computational resources, we process only the read alignments associated with filtered, highly-confidence cell barcodes. The output of this analysis includes the BAM files containing the alignments compatible with each feature category and some statistics on them. Specifically, we defined the following feature categories: $C_1$: the $3'$ terminal exon of transcripts, $C_2$: the $3'$ terminal kilobase of spliced transcripts, $C_3$: the $3'$ terminal kilobase of unspliced transcripts, $C_4$: the exon-exon junctions of spliced transcripts, $C_5$: the intron-exon and exon-intron junctions of unspliced transcripts, $C_6$: exons, and $C_7$: introns. Note that because features in different categories might share genomic loci, for example, $C_1$ is mostly a subset of $C_2$ and is a complete subset of $C_6$, read alignments can be assigned multiple categories. Among those categories, $C_1$, $C_2$, and $C_3$ represent the expected read mapping sites – the terminal kilobase of polyadenylated RNAs. Although most features in $C_1$ also belong to $C_2$ and $C_3$, except for those transcripts that have a terminal exon longer than $1,000$ bases, we define it as an explicit category because reads falling into $C_1$ are compatible with both spliced and unspliced transcripts and, therefore, have an ambiguous splicing status. On the contrary, $C_4$ and $C_5$ are the exclusive features in spliced and unspliced transcripts, respectively. Finally, $C_6$ and $C_7$ are used to take all other transcriptomic reads incompatible with the first five categories.

In the preparation step, we first filtered the read alignments associated with the high-confidence cell barcodes using the `samtools` program (Li et al., 2009). Those high-confidence cell barcodes were downloaded along with the datasets. Then, we split those alignments into two types, spliced alignments, and contiguous alignments, according to whether or not their CIGAR string includes a character "N".

Identifying alignments falling into categories $C_1$, $C_3$, $C_6$, and $C_7$ is straightforward. We used the `bedtools intersect` program (Quinlan and Hall, 2010) to find the contiguous alignments that are contained entirely within at least one feature in the corresponding feature category's BED file.

When finding alignments falling into category $C_5$, we represented each exon-intron junction as a six-base genomic interval centered at the junction and utilized the `bedtools intersect` program to find the contiguous alignments that contain at least one such interval.

Because `bedtools intersect` is unaware of genome-to-transcriptome conversion, we developed custom pipelines to identify alignments from categories $C_2$ and $C_4$, the categories related to splicing events. To find $C_4$ alignments, we filtered the spliced alignments that (a) are not contained entirely within any exon, (b) contain at least one exon's $5'$ end and one exon's $3'$ end, and (c) start and end in exons. The $5'$ end and $3'$ end of each exon are represented by the terminal three bases at the $5'$ and $3'$ end of the exon, respectively.

To find $C_2$ alignments, we filtered the splice and contiguous alignments that (a) are contained within the continuous genomic interval representing the terminal kilobase of spliced transcripts (including intronic regions) and (b) start and end in exons. The genomic intervals used in (a) are obtained by retracing the exonic bases of each spliced transcript on the genome from its $3'$ end until either reaching its $1,000$-th base or the $5'$ end. Notice that because the contiguous genomic intervals are extracted according to the accumulative length of exons, the retrieved genomic spans contain both the exons and the introns associated with the terminal kilobase of each spliced transcript and will be longer than $1,000$ bases if containing introns.

Then, we turned our attention to a large read population that was excluded in the above steps – the reads compatible with genes in the reverse complement, antisense orientation. Note that this analysis excluded all UMIs associated with sense reads, which were processed in the above steps. To be specific, we first found the antisense alignments by calling `bedtools intersect` with the `-S` flag. Then, we parsed those antisense alignments in a Python version 3.10.13 environment using `pysam`(Bonfield et al., 2021) to filter out the alignments from UMIs with sense alignments.

Finally, using a similar strategy, we extracted the intergenic alignments by calling `bedtools intersect` but with its `-v` flag to select only the reads that do not intersect with any transcriptomic regions in both forward and reverse complement orientations.

##### A.3 The statistical test for the abundance of intergenic reads near open chromatin regions

In this section, we describe the strategy used in this work to show the statistical significance of the enrichment of intergenic reads near intergenic open chromatin regions (OCRs). We define an intergenic region $I$, as the interval between a pair of adjacent genes. We define an intergenic OCR region $O_j$ as an OCR that does not intersect with any gene annotation. Moreover, we define a count function $c(\cdot,\cdot)$ and a size function $s(\cdot,\cdot)$. Both functions take two parameters. The first parameter is a set of intervals, and the second parameter is the flanking length that will be added to both ends of each interval when computing the count or size. Overlapping intervals after flanking will be concatenated before the calculation. $c(\cdot,\cdot)$ counts the UMIs compatible with the given intervals. $s(\cdot,\cdot)$ measures the size of the active intervals. Active intervals are the intervals that are associated with UMIs, i.e., have a non-zero count. The flanking length of intergenic regions is zero. The flanking length of OCRs is $4,000$.

For a chromosome $k$ in each cell, we first compute the total count and size of its intergenic regions $I$s and OCRs $O$s.

$$
\begin{aligned}
C_k^I &= c(\{I\}_k, 0) \\
C_k^O &= c(\{O\}_k, 4000) \\
S_k^I &= s(\{I\}_k, 0) \\
S_k^O &= s(\{O\}_k, 4000) \\
R_k^C &= \frac{C_k^I}{C_k^O} \\
R_k^S &= \frac{S_k^I}{S_k^O}
\end{aligned}
$$

where $\{I\}_k$ and $\{O\}_k$ represents the intergenic regions and OCRs on chromosome $k$, $C_k^I$ and $C_k^O$ are the total count of $\{I\}_k$ and $\{O\}_k$, $S_k^I$ and $S_k^O$ are the total size of active $\{I\}_k$ and $\{O\}_k$, $R_k^C$ is the ratio of $C_k^I$ and $C_k^O$, and $R_k^S$ is the ratio of $S_k^I$ and $S_k^O$

Next, we average over all cells to get $\bar{r}_C$ and $\bar{r}_S$, the average size and count ratio of intergenic regions and OCRs on each chromosome. Then, we test if $R_k^C$ and $R_k^S$ of all chromosomes follow the same distribution by applying a $t$-test. The null hypothesis is that, if intergenic reads are randomly distributed on the intergenic regions, the size and count ratio of intergenic regions and OCRs should follow the same distribution, so their ratio should follow a normal distribution with a mean of 1. Under the alternative hypothesis, we expect the mean of their ratio to be significantly different than 1.

#### Appendix B Supplementary Results

##### B.1 transcript composition analysis

In this analysis, we explored the composition of transcripts. From this analysis, we aim to build a comprehensive understanding of the introns and exons contained in each transcript and gene. We applied the analysis pipeline introduced in section A.1 to the standard gene annotations downloaded from 10X and *scRNA-seq optimized gene annotations* correspondingly. We find that the two sets of gene annotations for both mouse and human do not have significant differences in the analysis results. As the analysis results from mouse and human references followed the same patterns, here we use the standard human reference as the example to explain the findings. We first explored the exons in transcripts. We find that more than 47% of transcripts consist of a single exon, and 91.8% of transcripts have at most 10 exons. Notice that single-exon transcripts do not have intronic regions and therefore cannot have unspliced reads and UMIs. We also find that the terminal exon of most multi-exon transcripts is shorter than $1,000$ bases, indicating that in most cases, the reads generated from the end of the spliced and unspliced versions of transcripts are indistinguishable. Then, we analyzed the internal poly(A) tracts in spliced and unspliced versions of each transcript. In our analysis, we defined internal poly(A) tracts as the adenine-single nucleotide repeats (A-SNR) longer than bases. Furthermore, because of the possibility of cDNA-priming 10x (2021), in the following text, we used poly(A) as the sum of identified poly(A) and poly(T) tracts. We found that only less than 35% of exons contain internal poly(A) tracts. On the contrary, more than 24% of introns have at least one poly(A) tract, and most have poly(A) tracts in the $5'$ kilobase. If those $5'$ poly(A) tracts are primed by poly(T) primers, exonic reads might be generated from unspliced transcripts given that the cDNA fragment length is usually less than $1,000$ bases.

In conclusion, from this analysis, we conclude that poly(A) tracts are prevalent in the transcriptome, especially in the intronic regions of unspliced transcripts. Our results indicate that resolving the splicing status of reads compatible with exons, which are shared by the spliced and unspliced versions of transcripts, is not a trivial task, because of the prevalence of poly(A) tracts at the $5'$ end of introns, which can result in valid exonic reads, and the fact that the exons of each transcript are shared by its spliced and unspliced versions.

#### B.2 Read and UMI category analysis

In this section, we explain the analysis we did for classifying reads and UMIs into the seven categories introduced in section A.2. Specifically, we defined the following feature categories: $C_1$: the $3'$ terminal exon of transcripts, $C_2$: the $3'$ terminal kilobase of spliced transcripts, $C_3$: the $3'$ terminal kilobase of unspliced transcripts, $C_4$: the exon-exon junctions of spliced transcripts, $C_5$: the intron-exon and exon-intron junctions of unspliced transcripts, $C_6$: exons, and $C_7$: introns. Note that because features in different categories might share genomic loci, for example, $C_1$ is mostly a subset of $C_2$ and is a complete subset of $C_6$, each read alignment could be assigned into multiple categories. Among those, $C_1$, $C_2$, and $C_3$ represent the expected read mapping sites – the terminal kilobase of polyadenylated RNAs. Although $C_1$, the terminal exon of transcripts, is a subset of both $C_2$ and $C_3$ except for those transcripts that have a terminal exon longer than $1,000$ bases, we defined it as an explicit category. This is because reads falling into $C_1$ are compatible with both spliced and unspliced transcripts and, therefore, have an ambiguous splicing status. On the contrary, $C_4$ and $C_5$ are the exclusive features in spliced and unspliced transcripts, respectively. Finally, $C_6$ and $C_7$ are used to take all other transcriptomic reads incompatible with the first five categories. As shown in fig. 4a, in the read level, the two standing out category sets are (1) $C_7$ and (2) $(C_1, C_2, C_3, C_6)$. Being assigned $C_1$ means the reads were only compatible with introns, and, therefore, originated from unspliced transcripts. For the second set, $(C_1, C_2, C_3, C_6)$, it is clear that those reads were compatible with the terminal exon of transcripts, so they had an ambiguous splicing status. As for the definitive read categories for spliced transcripts, $C_4$ and $(C_2, C_4)$, they only account for 2.4% of reads. Similar results can be found for all processed datasets from both cell and nucleus samples. After grouping reads by their UMIs and applying the same strategy to find the category set of UMIs, as shown in fig. 4b, we find that the most outstanding groups are still the two representing ambiguous and spliced splicing statuses, agree with fig. 4a. This suggests that except for reads that are only compatible with $C_7$, the splicing status of 70% of the UMIs cannot be easily determined (Supplementary files). This finding highlights the necessity of designing sophisticated methods for resolving reads and UMIs' splicing status, as proposed in He et al. (2023). Interestingly, when using *scRNA-seq optimized gene annotations* (Pool et al., 2023), most of the intronic reads and UMIs were shifted to $(C_1, C_6, C_7)$, and were assigned an ambiguous splicing status (Supplementary files). The main reason was that the *spliceu* reference generated from *scRNA-seq optimized gene annotations* contained two conflicting sets of intronic regions. The intronic sequences contained in *scRNA-seq optimized gene annotations* were assigned a spliced status in *spliceu*. The intronic sequences created by *spliceu* were assigned an unspliced status in *spliceu*. Therefore, all intronic reads were assigned as ambiguous when using the *spliceu* generated from *scRNA-seq optimized gene annotations*. In this work, we focused on the results from the standard 10X gene annotations instead of *scRNA-seq optimized gene annotations*.

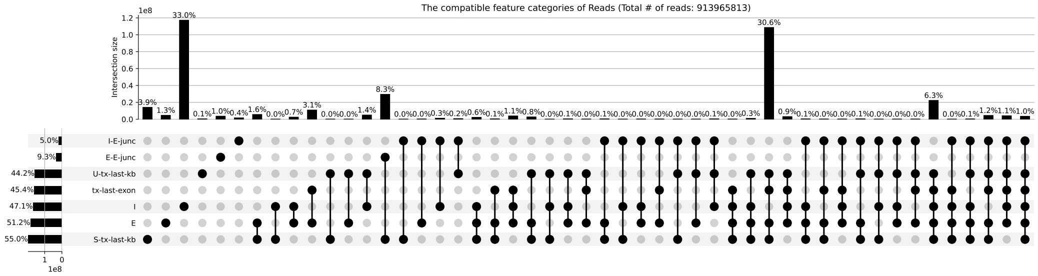


(a) Upset plot for reads in defined categories

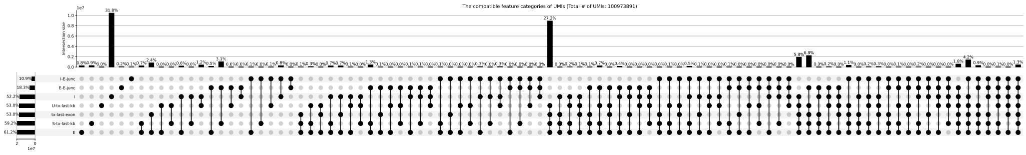

(b) Upset plot for UMIs in defined categories

**Fig. 4.** The genomic feature categories compatible with each (a) read and (b) UMI for the human PBMC scMultiome datasets from a nucleus sample. The dots and vertically connected dots represent the feature category and the set of categories compatible with a group of reads. The cardinality of each group of reads is shown on top of the dots in the histogram. In the plots. `tx-last-exon` means the $3'$ terminal exon of transcripts ($C_1$). The spliced and unspliced versions of each transcript share the same terminal exon. `S-tx-last-kb` means the $3'$ kilobase of spliced transcripts ($C_2$). `U-tx-last-kb` means the $3'$ kilobase of unspliced transcripts ($C_3$). `E-E-junc` means exon-exon junctions ($C_4$). `I-E-junc` stands for the intron-exon or exon-intron junctions ($C_5$). `E` stands for exons ($C_6$). `I` stands for introns ($C_7$).

#### Appendix C Supplementary figures and tables

In this section, we listed all additional figures and tables related to this work. To note that for some datasets, if the results were not applicable for generating figures or the analyses were too slow to be run, we excluded them from this section. For example, in the antisense read analysis, the mouse brain scMuiltiome datasets did not produce any differentially expressed genes due to the low number of antisense counts. We therefore excluded its DEG upset plots in this section. Furthermore, because some steps occasionally take a non-trivial amount of time to run for large datasets, for example, clustering cells into a specific number of clusters requires a clustering resolution tuning step. Because for large datasets, the individual iteration of this step might take very long to run, the entire tuning steps can take up to days to finish. Due to the limitation of computing resources, in this section, we only showed the jobs that have been done within the time limit of the University server. We also decided to display the antisense and intergenic analysis from *scRNA-seq optimized gene annotations* if the results from the standard 10X gene annotations were absent. This is because splicing status was not considered in these two types of analysis and we found that the corresponding analysis results and conclusions are consistent for these two types of gene annotations. If not explicitly marked, all figures and tables shown below are from the standard 10X gene annotations.

**Table 1.** In this table, we list the abbreviations used in this work.

| Abbreviation | Description |
|---|---|
| scRNA-seq | Single-cell RNA-sequencing |
| scATAC-seq | Single-cell sequencing assay for transposase-accessible chromatin |
| scMultiome | single-cell multiome ATAC+RNA assays |
| A-SNR | adenine-single nucleotide repeat |
| UMIs | unique molecule identifier |
| OCR | open chromatin regions |
| cCRE | candidate *cis*-regulatory element |
| *spliceu* | the *spliced+unspliced* augmented gene annotations |
| U | Unspliced splicing status |
| S | Spliced splicing status |
| A | Ambiguous splicing status |
| ARI | Adjusted Rand index |
| NMI | Normalized mutual information |
| FMI | Fowlkes–Mallows index |
| DEG | Differentially expressed gene |
| PBMC | Peripheral blood mononuclear cell |

**Table 2.** In this table, we list the annotation of all mathematical symbols used in this work.

| Symbol | Read analysis type | Description |
|---|---|---|
| $M_{\mathrm{S}}$ | Sense | The count matrix containing the sense gene counts of spliced splicing status |
| $M_{\mathrm{U}}$ | Sense | The count matrix containing the sense gene counts of unspliced splicing status |
| $M_{\mathrm{A}}$ | Sense | The count matrix including the sense gene count of ambiguous splicing status |
| $M_{\mathrm{SA}}$ | Sense | The count matrix including the sense gene count of spliced and ambiguous splicing status. This is the sum of $M_{\mathrm{S}}$ and $M_{\mathrm{A}}$. |
| $M_{\mathrm{USA}}$ | Sense | The count matrix including the sense gene counts of all splicing statuses. This is the sum of $M_{\mathrm{U}}$, $M_{\mathrm{S}}$, and $M_{\mathrm{A}}$. |
| $M_{\mathrm{std}}$ | Cell type annotation | The standard count matrix used for annotating cell types with `sctype`. For datasets generated from cell samples, this is $M_{\mathrm{SA}}$. For datasets generated from nucleus samples, this is $M_{\mathrm{USA}}$. |
| $M_{\mathrm{S}}^{\mathrm{rc}}$ | Antisense | The count matrix containing the antisense gene counts of spliced splicing status. |
| $M_{\mathrm{U}}^{\mathrm{rc}}$ | Antisense | The count matrix containing the antisense gene counts of unspliced splicing status. |
| $M_{\mathrm{A}}^{\mathrm{rc}}$ | Antisense | The count matrix including the antisense gene count of ambiguous splicing status. |
| $M_{\mathrm{anti}}$ | Antisense | The count matrix including the antisense gene counts of all splicing statuses. This is the sum of $M_{\mathrm{U}}^{\mathrm{rc}}$, $M_{\mathrm{S}}^{\mathrm{rc}}$, and $M_{\mathrm{A}}^{\mathrm{rc}}$. |
| $M_{\mathrm{sense}}$ | Antisense | Analog of $M_{\mathrm{USA}}$. The count matrix including the *sense* gene counts of all splicing statuses. |
| $M_{\mathrm{imputed}}$ | Antisense | The count matrix generated by imputing the zero entry values in $M_{\mathrm{sense}}$ as 1 if the corresponding entry values in $M_{\mathrm{anti}}$ is nonzero. |
| $M_{\mathrm{genic}}$ | Antisense | The count matrix including the gene counts of *all* splicing statuses from sense *and* antisense orientation. This is the sum of $M_{\mathrm{sense}}$ and $M_{\mathrm{anti}}$. |
| $F$ | Intergenic | A feature set including the genomic intervals of a genomic feature set. This can be either the ATAC-seq peaks discovered from a scATAC-seq experiment or the cCREs from the SCREEN project. |
| $F_{\mathrm{ig}}$ | Intergenic | A subset of the $F$ including only the features from intergenic regions. |
| $F_{\mathrm{nc}}$ | Intergenic | A subset of the $F$ including only the features from regions that are *not* related to protein-coding genes. |
| $M_{\mathrm{ig}}^{F_{\mathrm{ig}}}$ | Intergenic | The count matrix including the counts of features in $F_{\mathrm{ig}}$ generated using the reads from UMIs that *is not* associated with reads compatible with gene annotations. |
| $M_{\mathrm{nc}}^{F_{\mathrm{nc}}}$ | Intergenic | The count matrix including the counts of features in $F_{\mathrm{nc}}$ generated using the reads from UMIs that *is not* associated with reads compatible with protein-coding gene annotations. |
| $M_{\mathrm{all}}^{F}$ | Intergenic | The count matrix including the counts of features in $F$ generated using the all reads. |
| $M_{\mathrm{std}}$ | Intergenic | Same as the $M_{\mathrm{std}}$ used for cell type annotation. |
| $M_{\mathrm{ocr}}$ | Intergenic | Analog of $M_{\mathrm{ig}}^{F_{\mathrm{ig}}}$. |
| $M_{\mathrm{int}}$ | Intergenic | The count matrix including the counts of each intergenic region between a pair of adjacent genes generated using the reads from UMIs that *is not* associated with reads compatible with gene annotations. |

**Table 3.** Details of the selected single-cell datasets. Rows represent an attribute of the dataset, and columns represent the datasets. The counts listed in this table were based on the standard human and mouse reference sets downloaded from the 10x Genomics website (section 2.1). The same table w.r.t. the *scRNA-seq optimized gene annotations* can be found in table 4. In *Tissue type*, PBMC stands for peripheral blood mononuclear cells, BMMC stands for Primary Bone Marrow Mononuclear Cells, CHVZ stands for cortex, hippocampus, and ventricular zone. *Data Source* records the source where the dataset can be downloaded. 10X stands for the 10X Genomics Website, GEO stands for Gene Expression Omnibus. *Cell Count* lists the number of high-confidence cells filtered from the sequencing results. The column named Sample Type indicates the sample type used for generating the sequencing data, either isolated cells or nuclei. *Txome* used in the row names stands for Transcriptome. This genomic feature category takes reads compatible with the transcriptome in the forward orientation. In contrast, the *Antisense* category takes reads compatible with the transcriptome in the reverse complementary orientation. Notice that the reads from UMIs associating with *Txome* reads are excluded in other categories. *Spliced*, *Unspliced*, and *Ambiguous* represent the splicing statuses assigned by `alevin-fry`, where the *Spliced* status is assigned to UMIs that are only compatible with spliced transcripts, the *Unspliced* status is assigned to UMIs that only compatible with unspliced transcripts, and the *ambiguous* status is assigned to UMIs that compatible with both spliced and unspliced transcripts. *cCREs* stands for candidate *cis*-regulatory elements Consortium. et al. (2020). *OCRs* stands for open chromatin regions. In this table, ATAC-seq peaks from the ATAC-seq modality in the multiomics datasets were used to represent the *OCRs* for all datasets of the same species and tissue type. In table 5 we showed the *OCR* counts when using the peaks from independent ATAC-seq datasets as the *OCR*.

| Attribute \ Name | pbmc_multiome | pbmc | bmmc_multiome | bmmc | human_brain_multiome | mouse_brain_multiome | mouse_brain |
|---|---|---|---|---|---|---|---|
| Data Source | 10X | 10X | GEO | GEO | 10X | 10X | 10X |
| Species | Human | Human | Human | Human | Human | Mouse | Mouse |
| Tissue Type | PBMC | PBMC | BMMC | BMMC | Cerebellum | CHVZ | CHVZ |
| Sample Type | Nuclei | Cells | Nuclei | Cells | Nuclei | Nuclei | Nuclei |
| Assay Type | RNA+ATAC | RNA | RNA+ATAC | RNA | RNA+ATAC | RNA+ATAC | RNA |
| Cell Count | 10, 974 | 11, 996 | 16, 360 | 5, 122 | 3, 233 | 4, 878 | 5, 973 |
| Development Stage | Adult | Adult | Adult | Adult | Adult | E18 | E18 |
| Txome Count | 35,455,902 | 145,584,599 | 26,012,597 | 60,402,509 | 36,902,650 | 50,144,228 | 37,354,772 |
| % Spliced Txome | 31.21% | 29.11% | 11.79% | 37.43% | 3.47% | 19.59% | 21.78% |
| % Unspliced Txome | 35.08% | 33.43% | 61.38% | 26.23% | 77.52% | 48.81% | 44.87% |
| % Ambiguous Txome | 33.7% | 37.46% | 26.83% | 36.34% | 19.01% | 31.61% | 33.35% |
| Antisense Count | 8,402,601 | 19,965,710 | 10,763,603 | 8,223,895 | 5,103,759 | 11,699,000 | 13,817,752 |
| % Spliced Antisense | 2.73% | 1.78% | 4.34% | 1.59% | 0.77% | 3.01% | 1.73% |
| % Unspliced Antisense | 84.76% | 85.37% | 76.02% | 85.67% | 89.32% | 83.27% | 88.23% |
| % Ambiguous Antisense | 12.51% | 12.85% | 19.64% | 12.74% | 9.91% | 13.72% | 10.04% |
| Intergenic Count | 4,359,845 | 11,091,646 | 5,344,171 | 5,173,453 | 4,740,010 | 6,094,866 | 2,722,559 |
| cCREs Intergenic Count | 2,394,025 | 6,321,381 | 3,060,628 | 2,477,817 | 2,926,915 | 1,896,725 | 1,093,156 |
| % cCREs Intergenic | 54.91% | 56.99% | 57.27% | 47.89% | 61.75% | 31.12% | 40.15% |
| cCREs Non-Coding Count | 5,036,044 | 14,540,471 | 5,610,987 | 5,808,574 | 6,230,140 | 3,980,877 | 3,191,966 |
| cCREs All Count | 38,010,260 | 158,482,068 | 33,741,606 | 64,314,069 | 38,189,293 | 40,786,052 | 35,387,627 |
| OCRs Intergenic Count | 1,645,032 | 3,576,295 | 2,222,147 | 1,840,623 | 1,257,817 | 1,801,766 | 943,361 |
| % OCRs Intergenic | 37.73% | 32.24% | 41.58% | 35.58% | 26.54% | 29.56% | 34.65% |
| OCRs Non-Coding Count | 3,949,744 | 10,313,740 | 4,724,716 | 4,866,989 | 3,433,407 | 4,049,269 | 3,163,351 |
| OCR All Count | 28,165,222 | 113,703,447 | 24,971,437 | 52,722,330 | 22,123,440 | 38,896,532 | 32,900,172 |

**Table 4.** Details of the selected single-cell datasets. This table shows the sample information as in table 3, but the counts were based on the *scRNA-seq optimized gene annotations* (section 2.1).

| Attribute \ Name | pbmc_multiome | pbmc | bmmc_multiome | bmmc | human_brain_multiome | mouse_brain_multiome | mouse_brain |
|---|---|---|---|---|---|---|---|
| Data Source | 10X | 10X | GEO | GEO | 10X | 10X | 10X |
| Species | Human | Human | Human | Human | Human | Mouse | Mouse |
| Tissue Type | PBMC | PBMC | BMMC | BMMC | Cerebellum | CHVZ | CHVZ |
| Sample Type | Nuclei | Cells | Nuclei | Cells | Nuclei | Nuclei | Nuclei |
| Assay Type | RNA+ATAC | RNA | RNA+ATAC | RNA | RNA+ATAC | RNA+ATAC | RNA |
| Cell Count | 10, 974 | 11, 996 | 16, 360 | 5, 122 | 3, 233 | 4, 878 | 5, 973 |
| Development Stage | Adult | Adult | Adult | Adult | Adult | E18 | E18 |
| Txome Count | 35,353,341 | 144,101,363 | 26,095,341 | 59,853,860 | 36,989,036 | 49,840,738 | 37,059,671 |
| % Spliced Txome | 49.99% | 49.02% | 36.06% | 54.99% | 30.3% | 55.65% | 58.45% |
| % Unspliced Txome | 6.71% | 6.48% | 11.52% | 5.06% | 17.95% | 6.27% | 5.59% |
| % Ambiguous Txome | 43.3% | 44.5% | 52.42% | 39.95% | 51.74% | 38.08% | 35.96% |
| Antisense Count | 8,496,602 | 20,153,098 | 10,871,110 | 8,304,741 | 5,153,998 | 11,727,145 | 13,855,665 |
| % Spliced Antisense | 30.26% | 30.46% | 31.54% | 30.43% | 29.96% | 42.89% | 43.3% |
| % Unspliced Antisense | 15.6% | 15.66% | 13.43% | 15.72% | 18.62% | 9.4% | 9.69% |
| % Ambiguous Antisense | 54.14% | 53.88% | 55.03% | 53.84% | 51.42% | 47.71% | 47.01% |
| Intergenic Count | 4,447,923 | 11,410,054 | 5,428,639 | 5,264,384 | 4,910,827 | 6,172,168 | 2,776,047 |
| cCREs Intergenic Count | 2,468,157 | 6,565,987 | 3,124,323 | 2,546,690 | 3,048,014 | 1,935,938 | 1,125,565 |
| % cCREs Intergenic | 55.49% | 57.55% | 57.55% | 48.38% | 62.07% | 31.37% | 40.55% |
| cCREs Non-Coding Count | 5,107,380 | 14,787,613 | 5,685,175 | 5,887,612 | 6,355,757 | 4,022,321 | 3,227,152 |
| cCREs All Count | 38,010,260 | 158,482,068 | 33,741,606 | 64,314,069 | 38,189,293 | 40,786,052 | 35,387,627 |
| OCRs Intergenic Count | 1,689,967 | 3,715,689 | 2,260,479 | 1,887,377 | 1,308,435 | 1,835,115 | 969,687 |
| % OCRs Intergenic | 37.99% | 32.57% | 41.64% | 35.85% | 26.64% | 29.73% | 34.93% |
| OCRs Non-Coding Count | 3,992,373 | 10,460,977 | 4,776,088 | 4,923,085 | 3,485,696 | 4,091,295 | 3,196,964 |
| OCR All Count | 28,165,222 | 113,703,447 | 24,971,437 | 52,722,330 | 22,123,440 | 38,896,532 | 32,900,172 |

**Table 5.** The Open Chromatin Regions(OCR)-associated UMI counts using independent ATAC-seq datasets. The human PBMC ATAC-seq dataset was downloaded from https://www.10xgenomics.com/resources/datasets/10k-human-pbmcs-atac-v2-chromium-x-2-standard. The mouse brain ATAC-seq dataset was downloaded from https://www.10xgenomics.com/resources/datasets/fresh-cortex-hippocampus-and-ventricular-zone-from-embryonic-mouse-brain-e-18-1-standard-1-1-0. The only difference between this table and the OCR-related counts in (Table 3) is that the OCRs used in (Table 3) came from the ATAC-seq peaks observed from the multi-omics datasets used in this work, while the OCRs used to generate the numbers in this table came from independent scATAC-seq datasets. The numbers shown here are consistent with the numbers shown in (Table 3), suggesting that the conclusions in this work hold for unpaired single-cell RNA+ATAC sequencing data.

| | pbmc_multiome | pbmc | mouse_brain_multiome | mouse_brain |
|---|---|---|---|---|
| OCRs Intergenic Count | 1,916,535 | 4,441,120 | 591,897 | 329,813 |
| % OCRs Intergenic | 43.96% | 40.04% | 9.71% | 12.11% |
| OCRs Non-Coding Count | 4,338,278 | 11,529,800 | 895,720 | 579,177 |
| OCR All Count | 31,825,314 | 129,767,568 | 14,222,887 | 11,992,065 |
| cCREs Intergenic Count | 2,394,025 | 6,321,381 | 1,896,725 | 1,093,156 |
| % cCREs Intergenic | 54.91% | 56.99% | 31.12% | 40.15% |
| cCREs Non-Coding Count | 5,036,044 | 14,540,471 | 3,980,877 | 3,191,966 |
| cCREs All Count | 38,010,260 | 158,482,068 | 40,786,052 | 35,387,627 |

**Table 6.** The Open Chromatin Regions(OCR)-associated UMI counts using independent ATAC-seq datasets. This table shows similar information as in table 5 but the counts were generated from the *scRNA-seq optimized gene annotations*.

| | pbmc_multiome | pbmc | mouse_brain_multiome | mouse_brain |
|---|---|---|---|---|
| OCRs Intergenic Count | 1,967,714 | 4,603,731 | 597,054 | 332,814 |
| % OCRs Intergenic | 44.24% | 40.35% | 9.67% | 11.99% |
| OCRs Non-Coding Count | 4,387,920 | 11,704,653 | 915,334 | 597,019 |
| OCR All Count | 31,825,314 | 129,767,568 | 14,222,887 | 11,992,065 |
| cCREs Intergenic Count | 2,468,157 | 6,565,987 | 1,935,938 | 1,125,565 |
| % cCREs Intergenic | 55.49% | 57.55% | 31.37% | 40.55% |
| cCREs Non-Coding Count | 5,107,380 | 14,787,613 | 4,022,321 | 3,227,152 |
| cCREs All Count | 38,010,260 | 158,482,068 | 40,786,052 | 35,387,627 |

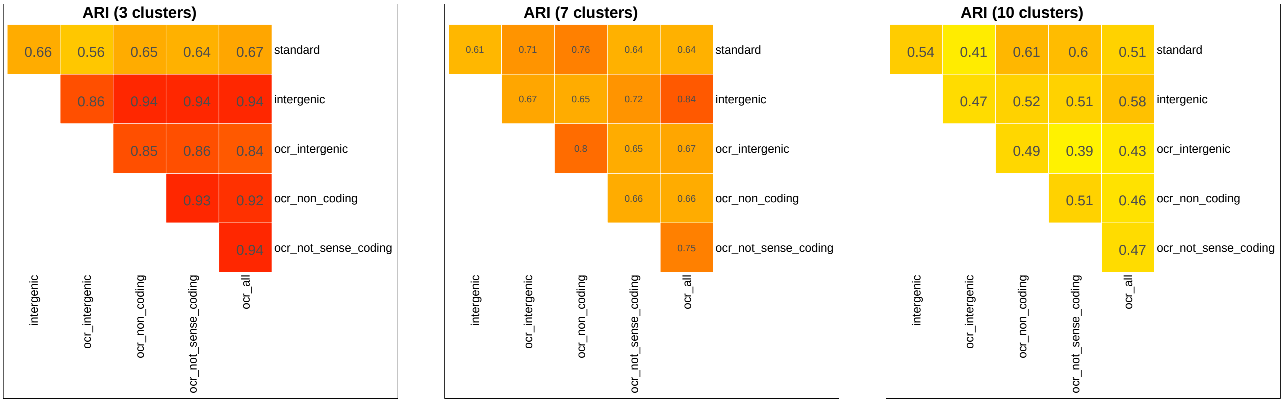


(a) (b) (c)

**Fig. 5.** The ARI of the clustering results generated from the *OCR* count matrices explored in section 2.6 for the human PBMC scMultiome dataset. `S`: spliced counts. `U`: unspliced counts. `A`: ambiguous counts. `SA`: spliced and ambiguous total counts. `USA`: spliced, unspliced, and ambiguous total counts. (a) represents the ARIs between the clusters discovered from different modalities using a low clustering resolution. (b) represents the results from a middle clustering resolution. (c) represents the results from a high clustering resolution.

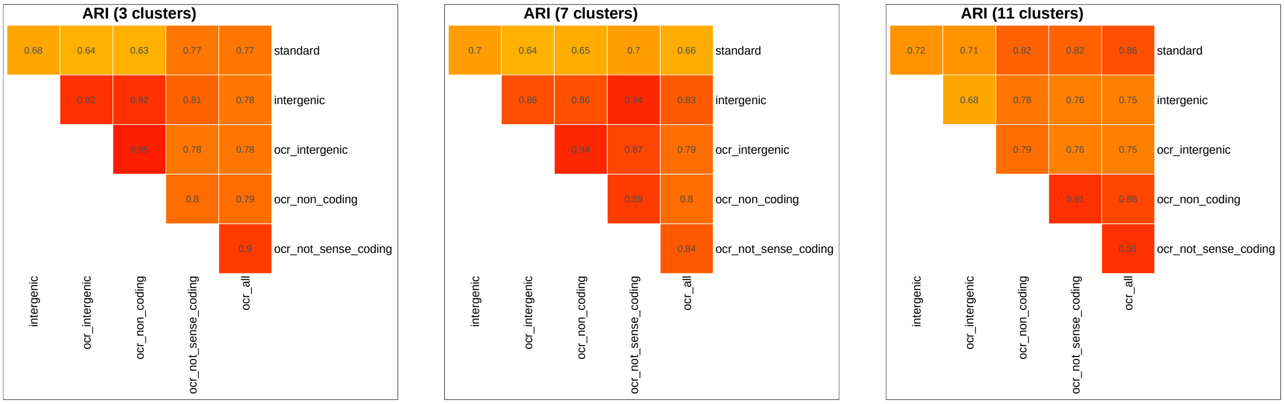


(a) (b) (c)

**Fig. 6.** The ARI of the clustering results generated from the *OCR* count matrices explored in section 2.6 for the human PBMC scRNA-seq dataset. `S`: spliced counts. `U`: unspliced counts. `A`: ambiguous counts. `SA`: spliced and ambiguous total counts. `USA`: spliced, unspliced and ambiguous total counts. (a) represents the ARIs between the clusters discovered from different modalities using a low clustering resolution. (b) represents the results from a middle clustering resolution. (c) represents the results from a high clustering resolution.

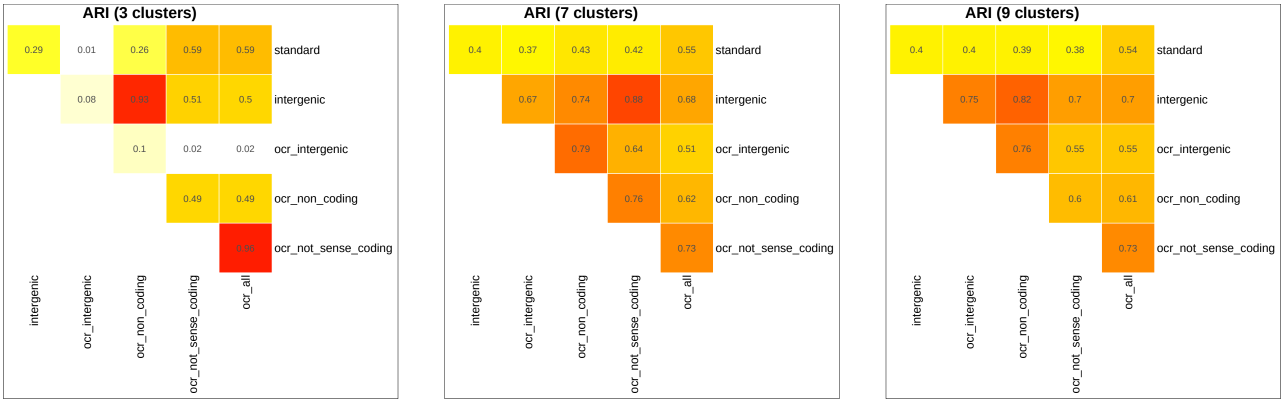


(a) (b) (c)

**Fig. 7.** The ARI of the clustering results generated from the *OCR* count matrices explored in section 2.6 for the human BMMC scMultiome dataset. `S`: spliced counts. `U`: unspliced counts. `A`: ambiguous counts. `SA`: spliced and ambiguous total counts. `USA`: spliced, unspliced, and ambiguous total counts. (a) represents the ARIs between the clusters discovered from different modalities using a low clustering resolution. (b) represents the results from a middle clustering resolution. (c) represents the results from a high clustering resolution.

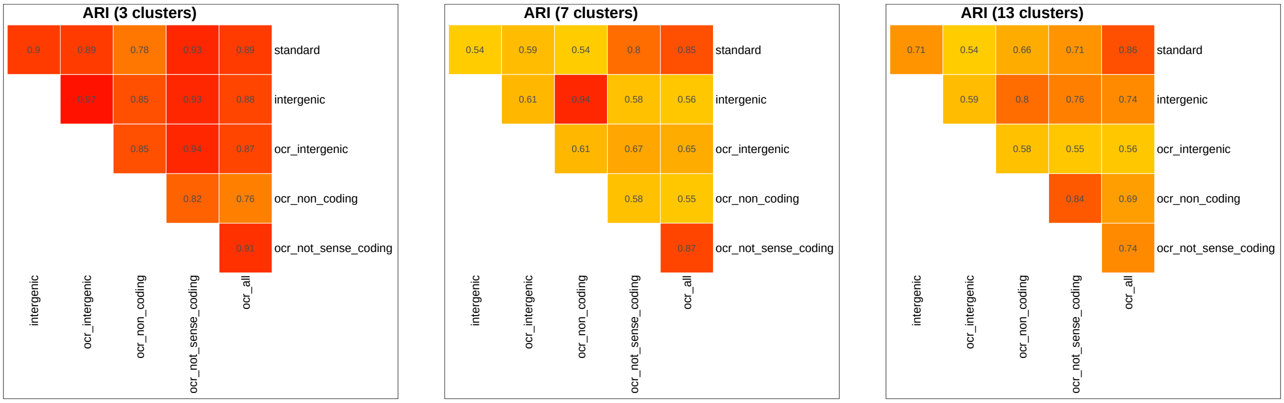


(a) (b) (c)

**Fig. 8.** The ARI of the clustering results generated from the *OCR* count matrices explored in section 2.6 for the human BMMC dataset. `S`: spliced counts. `U`: unspliced counts. `A`: ambiguous counts. `SA`: spliced and ambiguous total counts. `USA`: spliced, unspliced, and ambiguous total counts. (a) represents the ARIs between the clusters discovered from different modalities using a low clustering resolution. (b) represents the results from a middle clustering resolution. (c) represents the results from a high clustering resolution.

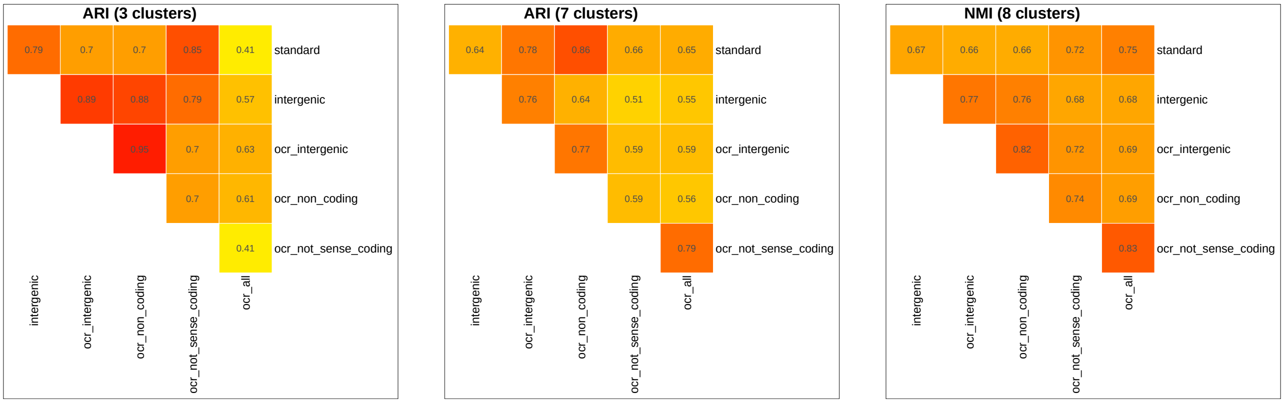


(a) (b) (c)

**Fig. 9.** The ARI of the clustering results generated from the *OCR* count matrices explored in section 2.6 for the human brain scMultiome dataset. `S`: spliced counts. `U`: unspliced counts. `A`: ambiguous counts. `SA`: spliced and ambiguous total counts. `USA`: spliced, unspliced, and ambiguous total counts. (a) represents the ARIs between the clusters discovered from different modalities using a low clustering resolution. (b) represents the results from a middle clustering resolution. (c) represents the results from a high clustering resolution.

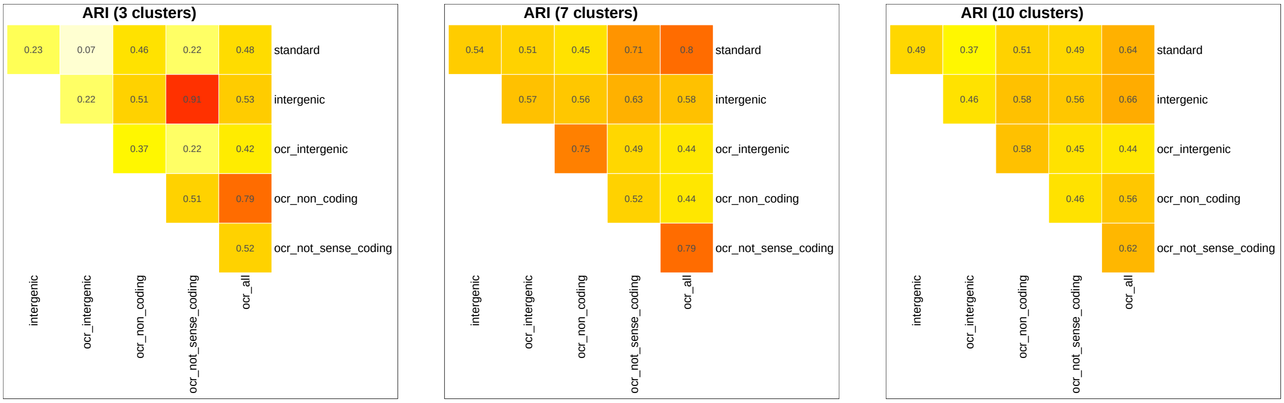


(a) (b) (c)

**Fig. 10.** The ARI of the clustering results generated from the *OCR* count matrices explored in section 2.6 for the mouse brain scMultiome dataset. `S`: spliced counts. `U`: unspliced counts. `A`: ambiguous counts. `SA`: spliced and ambiguous total counts. `USA`: spliced, unspliced, and ambiguous total counts. (a) represents the ARIs between the clusters discovered from different modalities using a low clustering resolution. (b) represents the results from a middle clustering resolution. (c) represents the results from a high clustering resolution.

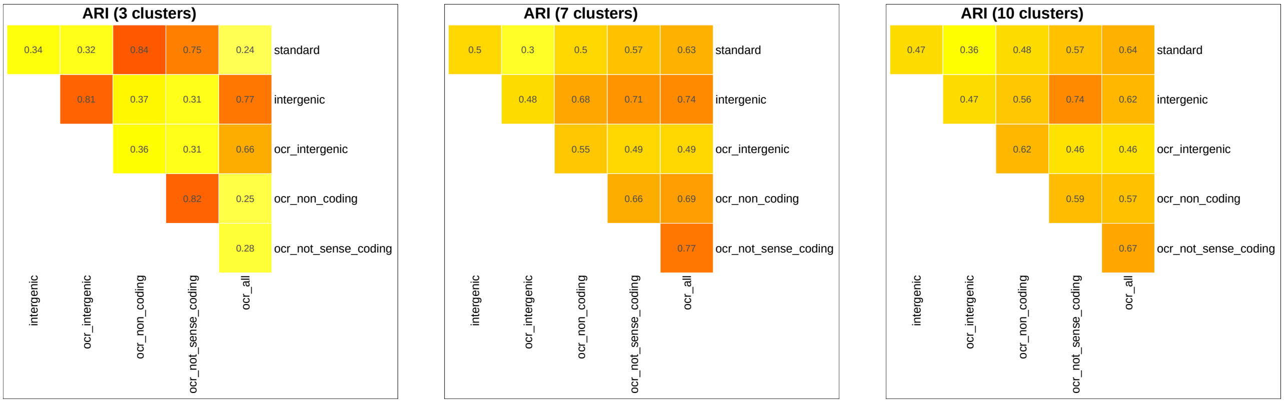


(a) (b) (c)

**Fig. 11.** The ARI of the clustering results generated from the *OCR* count matrices explored in section 2.6 for the mouse brain dataset. `S`: spliced counts. `U`: unspliced counts. `A`: ambiguous counts. `SA`: spliced and ambiguous total counts. `USA`: spliced, unspliced, and ambiguous total counts. (a) represents the ARIs between the clusters discovered from different modalities using a low clustering resolution. (b) represents the results from a middle clustering resolution. (c) represents the results from a high clustering resolution.

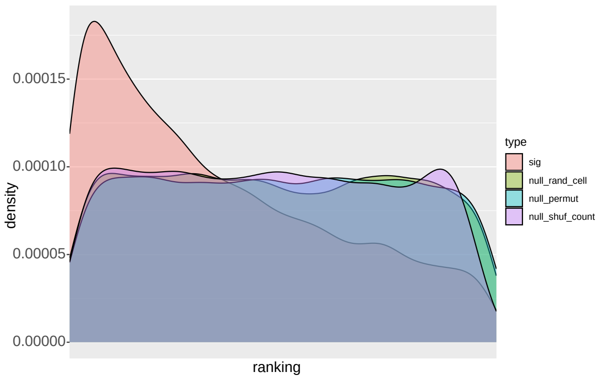


(a)

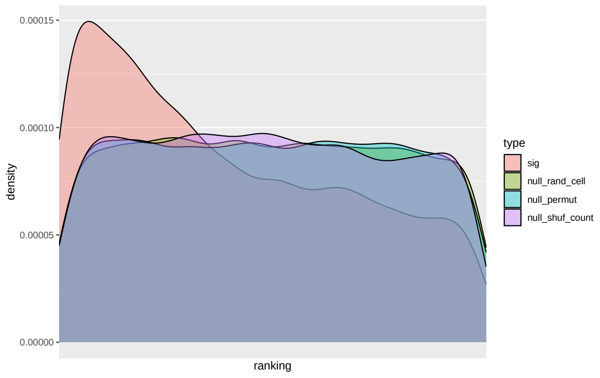


(b)

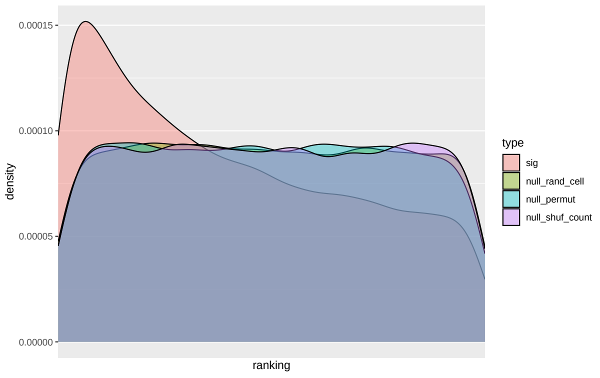


(c)

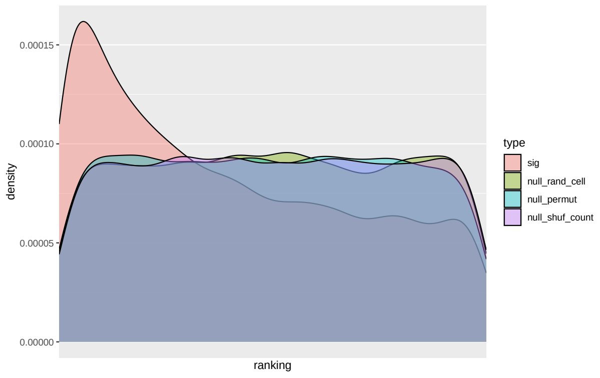


(d)

**Fig. 12.** The ranking of the cosine similarity of cell's ATAC-seq peak counts to its *iOCR* counts among all cells (section 2.6) for the human PBMC scMultiome dataset. The other three curves represent the three null distributions. `sig`: The ranking distribution to be tested. `null_rand_cell`: The null distribution is generated by randomly selecting a cell as the target to compare with. `null_permut`: The null distribution is generated by permuting the ranking directly instead of ranking the cosine similarities. `null_shuf_count`: The null distribution is generated by shuffling the count matrices before computing the cosine similarities. The four plots show the four sets of OCRs used for generating the count matrices (section 2.6). (a) Intergenic OCRs. (b) OCRs from intergenic regions and non-protein coding genes' regions. (c) OCRs from intergenic regions, non-protein coding genes' regions, and the reverse complement strand of protein-coding genes. (d) All OCRs.

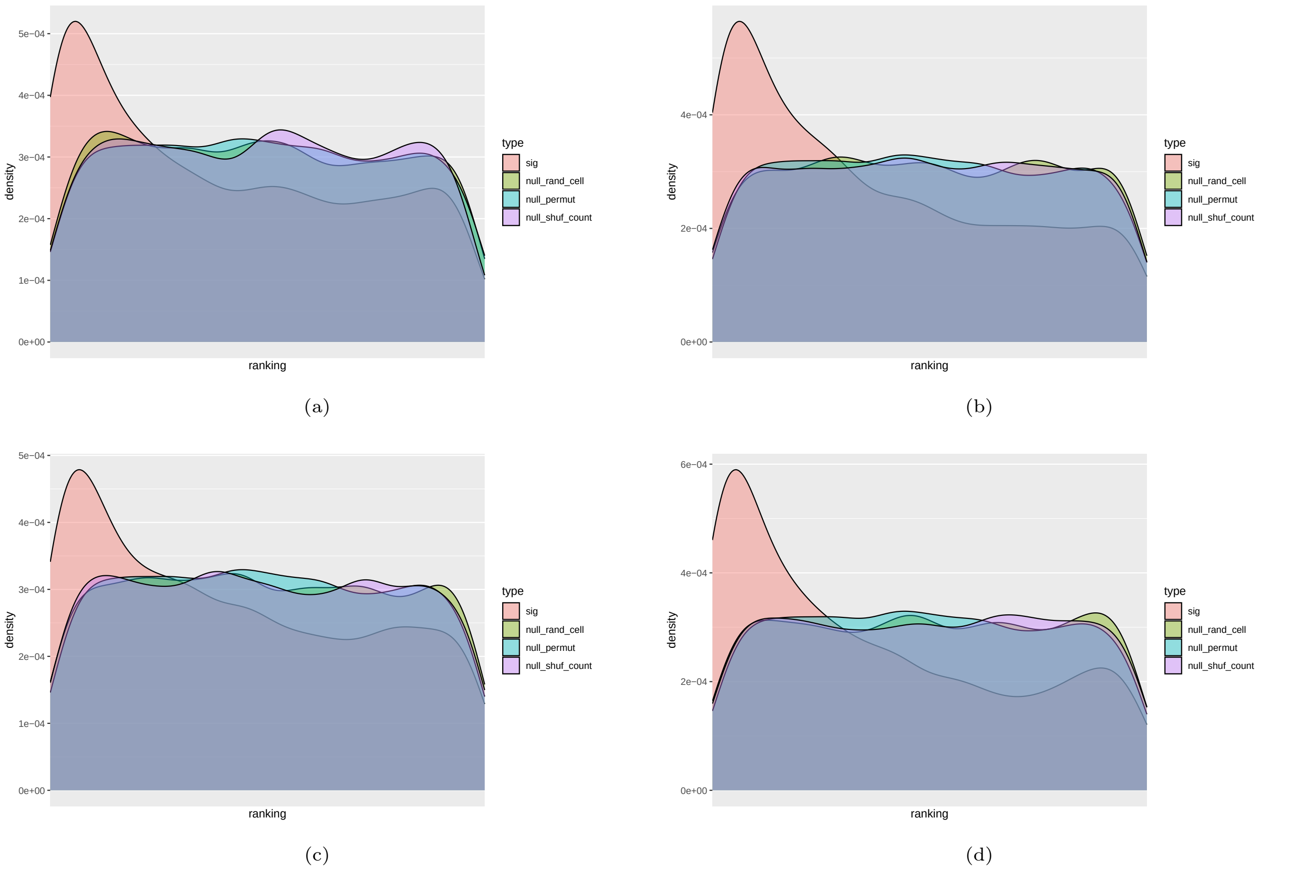


**Fig. 13.** The ranking of the cosine similarity of cell's ATAC-seq peak counts to its *iOCR* counts among all cells (section 2.6) for the mouse brain scMultiome dataset. The other three curves represent the three null distributions. `sig`: The ranking distribution to be tested. `null_rand_cell`: The null distribution is generated by randomly selecting a cell as the target to compare with. `null_permut`: The null distribution is generated by permuting the ranking directly instead of ranking the cosine similarities. `null_shuf_count`: The null distribution is generated by shuffling the count matrices before computing the cosine similarities. The four plots show the four sets of OCRs used for generating the count matrices (section 2.6). (a) Intergenic OCRs. (b) OCRs from intergenic regions and non-protein coding genes' regions. (c) OCRs from intergenic regions, non-protein coding genes' regions, and the reverse complement strand of protein-coding genes. (d) All OCRs.

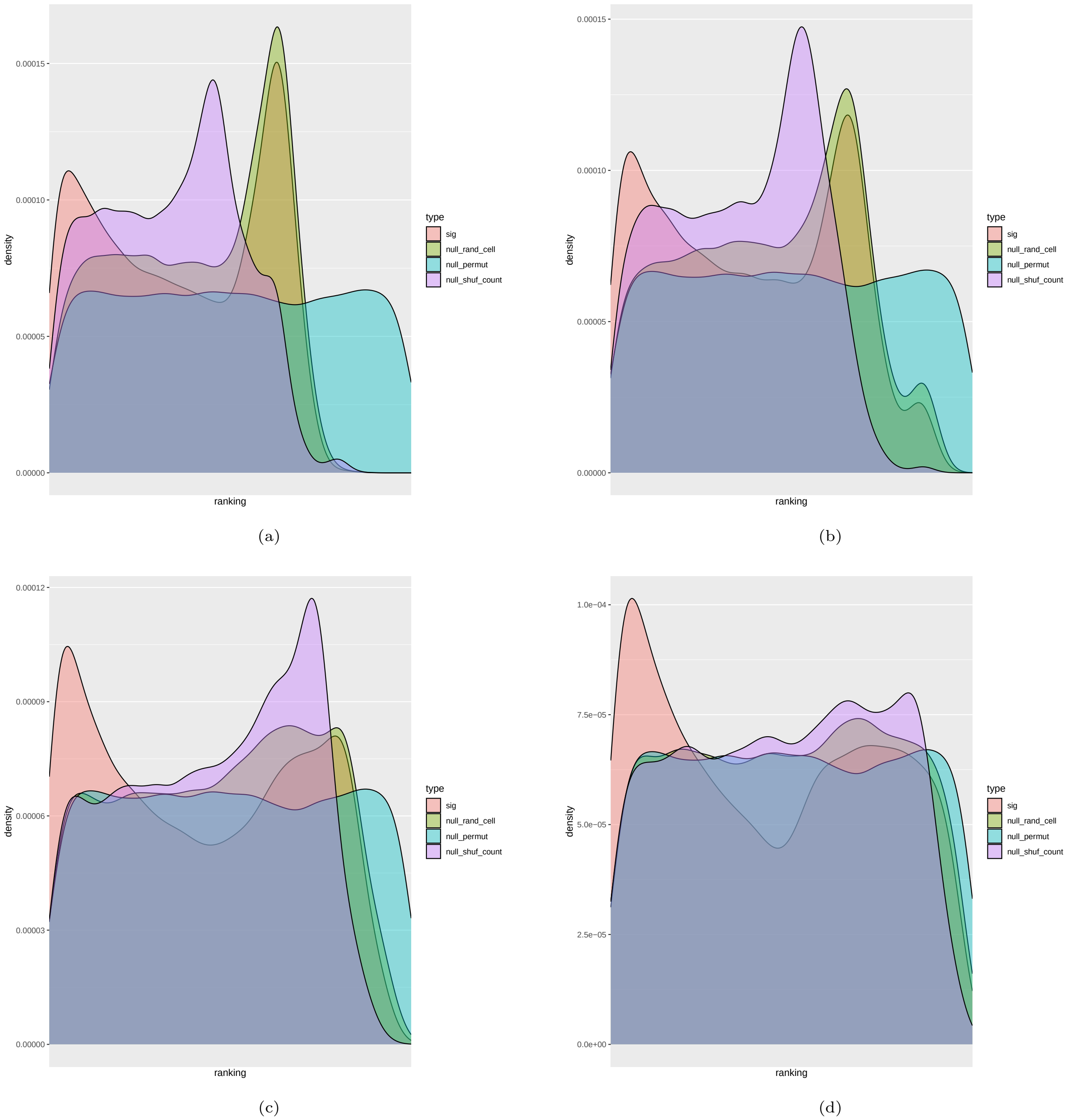


**Fig. 14.** The ranking of the cosine similarity of cell's ATAC-seq peak counts to its *iOCR* counts among all cells (section 2.6) for the human BMMC scMultiome dataset. The other three curves represent the three null distributions. `sig`: The ranking distribution to be tested. `null_rand_cell`: The null distribution is generated by randomly selecting a cell as the target to compare with. `null_permut`: The null distribution is generated by permuting the ranking directly instead of ranking the cosine similarities. `null_shuf_count`: The null distribution is generated by shuffling the count matrices before computing the cosine similarities. The four plots show the four sets of OCRs used for generating the count matrices (section 2.6). (a) Intergenic OCRs. (b) OCRs from intergenic regions and non-protein coding genes' regions. (c) OCRs from intergenic regions, non-protein coding genes' regions, and the reverse complement strand of protein-coding genes. (d) All OCRs. The binomial distributions for all but `null_permut` were caused by the sparsity of the *OCR* count matrices. Because most cells had only a few *OCR* counts, huge ties were observed in the ranking distribution. These ties disturbed the ranking distribution and led to binomial distribution-like curves. As `null_permut` was generated by permitting samples directly, the sparsity could not affect its distribution.

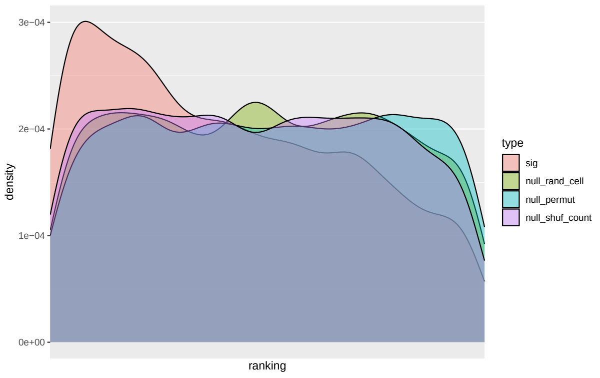


(a)

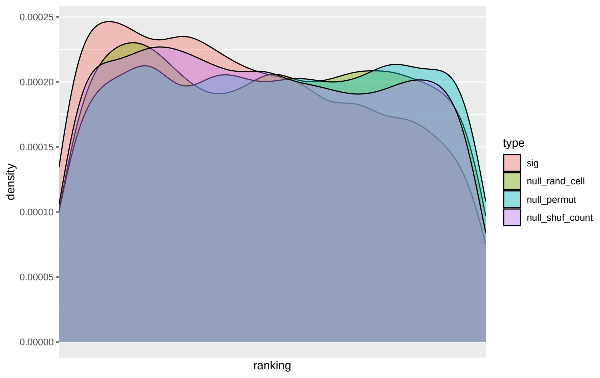


(b)

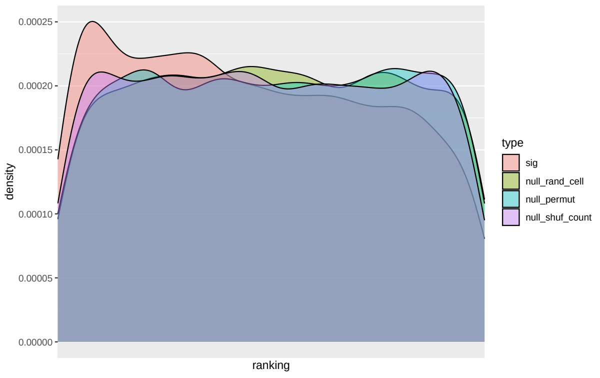


(c)

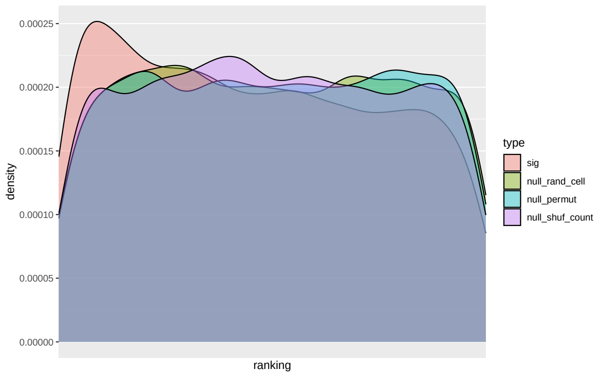


(d)

**Fig. 15.** The ranking of the cosine similarity of cell's ATAC-seq peak counts to its *iOCR* counts among all cells (section 2.6) for the human brain scMultiome dataset. The other three curves represent the three null distributions. `sig`: The ranking distribution to be tested. `null_rand_cell`: The null distribution is generated by randomly selecting a cell as the target to compare with. `null_permut`: The null distribution is generated by permuting the ranking directly instead of ranking the cosine similarities. `null_shuf_count`: The null distribution is generated by shuffling the count matrices before computing the cosine similarities. The four plots show the four sets of OCRs used for generating the count matrices (section 2.6). (a) Intergenic OCRs. (b) OCRs from intergenic regions and non-protein coding genes' regions. (c) OCRs from intergenic regions, non-protein coding genes' regions, and the reverse complement strand of protein-coding genes. (d) All OCRs.

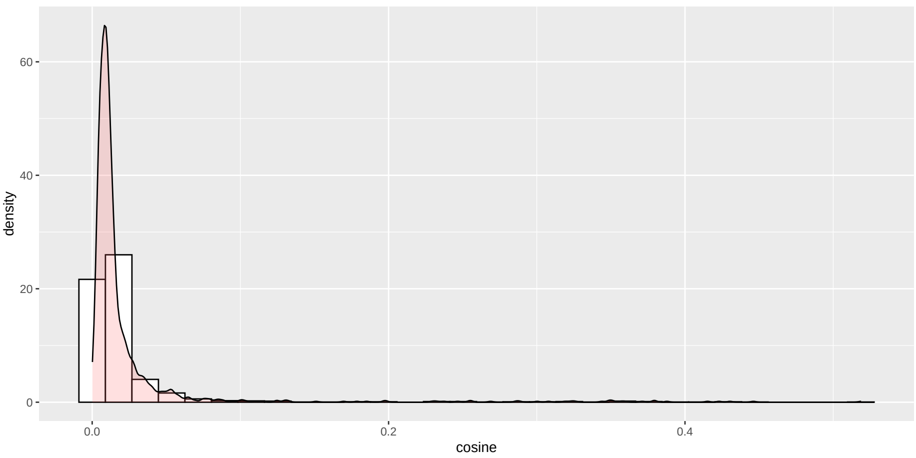


(a)

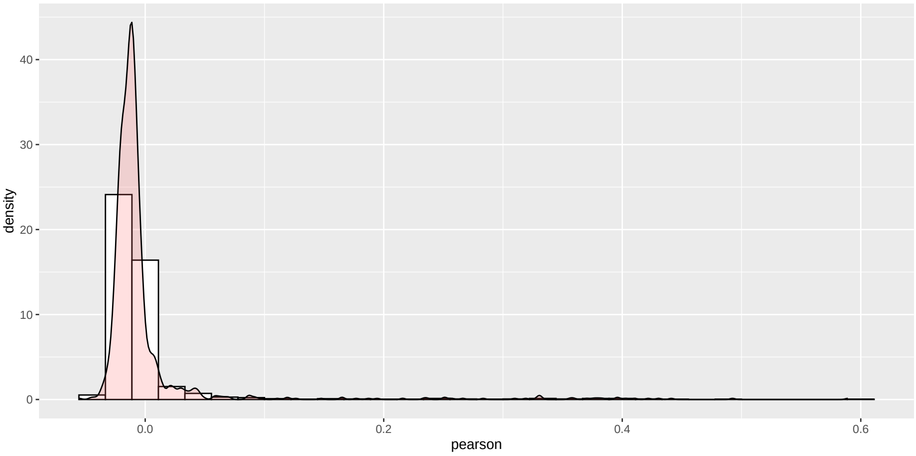


(b)

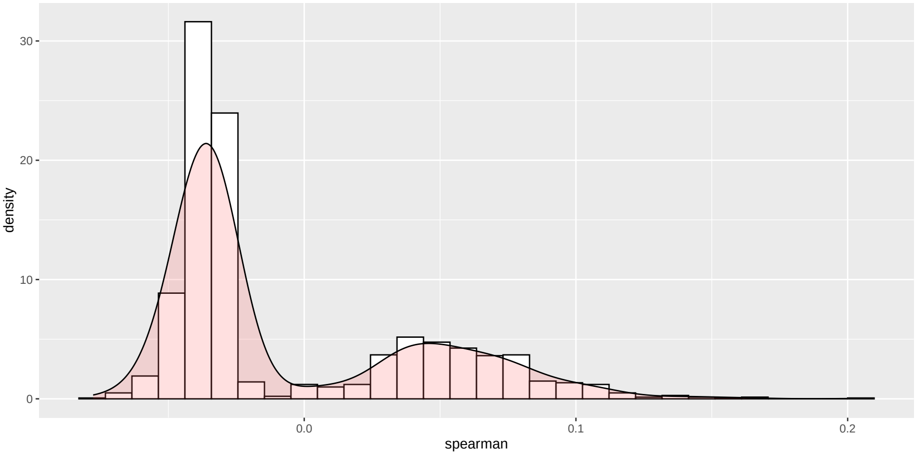


(c)

**Fig. 16.** An example of the (a) cosine similarity (b) Pearson correlation coefficient and (c) Spearman correlation coefficient of cells' sense and antisense counts using a shuffled sense and antisense count matrix. The shuffled sense and antisense count matrices were generated by randomly shuffling the entries in the sense and antisense count matrix separately.

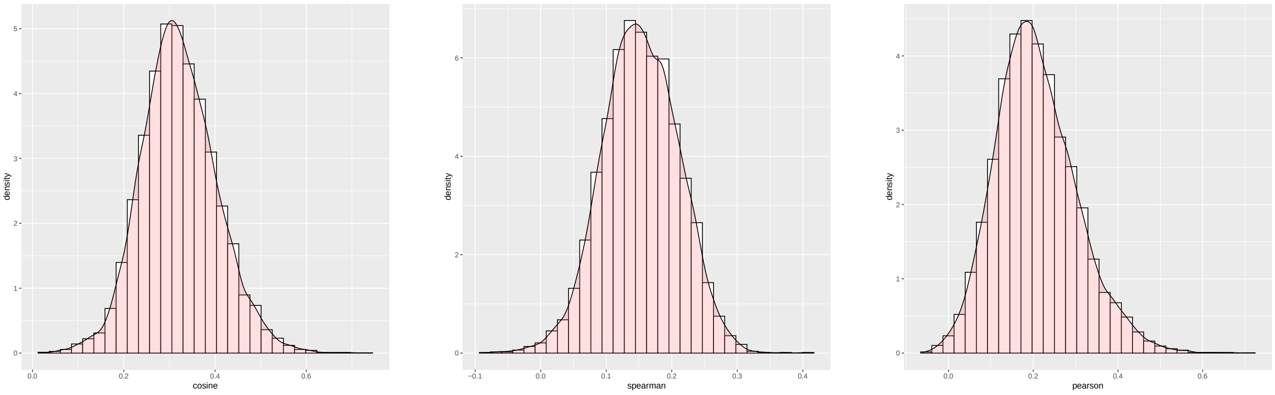


(a) (b) (c)

**Fig. 17.** The similarity scores of the sense and antisense counts of cells explored in section 2.5 for the human PBMC scMultiome dataset. (a): cosine similarity. (b): Spearman correlation coefficient ($\rho$). (c): Pearson correlation coefficient ($r$).

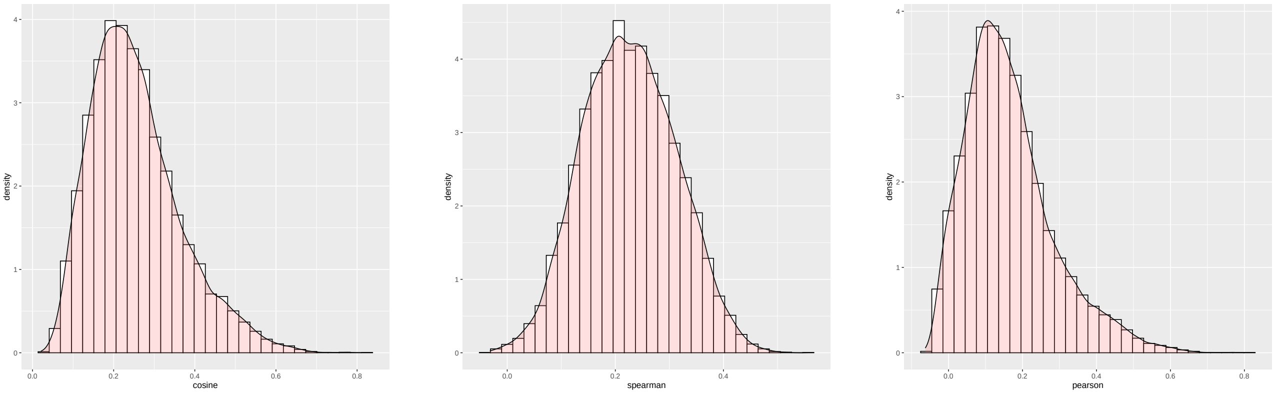


(a) (b) (c)

**Fig. 18.** The similarity scores of the sense and antisense counts of cells explored in section 2.5 for the human PBMC dataset. (a): cosine similarity. (b): Spearman correlation coefficient ($\rho$). (c): Pearson correlation coefficient ($r$).

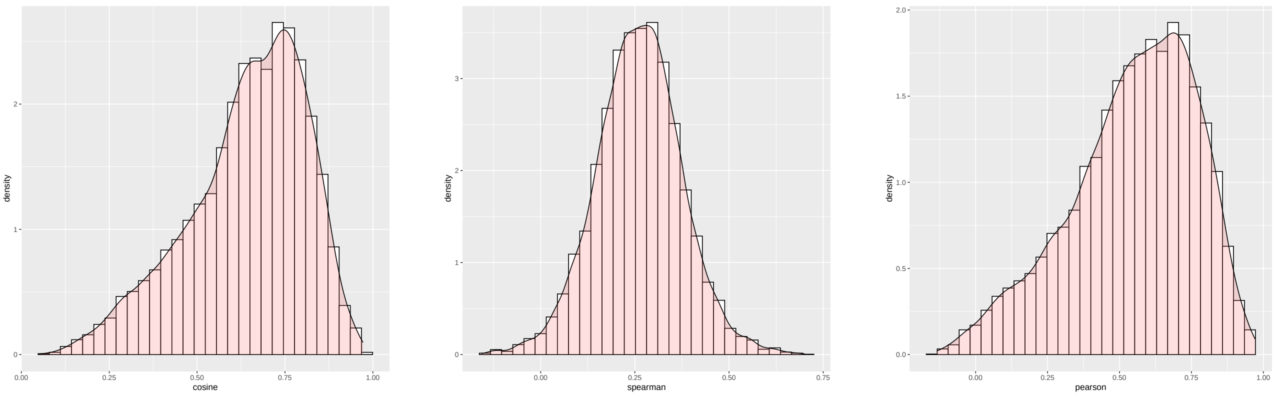


(a) (b) (c)

**Fig. 19.** The similarity scores of the sense and antisense counts of cells explored in section 2.5 for the human BMMC scMultiome dataset. (a): cosine similarity. (b): Spearman correlation coefficient ($\rho$). (c): Pearson correlation coefficient ($r$).

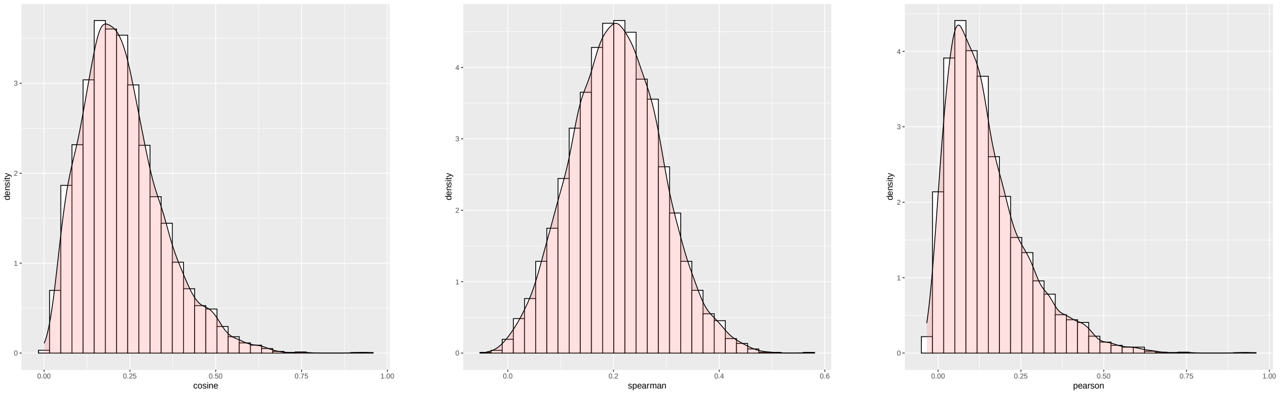


(a) (b) (c)

**Fig. 20.** The similarity scores of the sense and antisense counts of cells explored in section 2.5 for the human BMMC dataset. (a): cosine similarity. (b): Spearman correlation coefficient ($\rho$). (c): Pearson correlation coefficient ($r$).

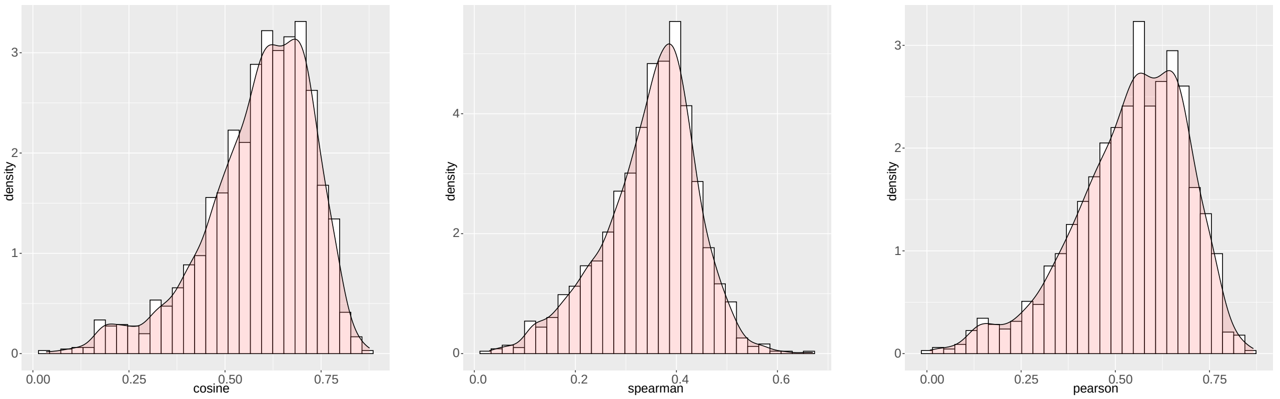


(a) (b) (c)

**Fig. 21.** The similarity scores of the sense and antisense counts of cells explored in section 2.5 for the human brain scMultiome dataset. (a): cosine similarity. (b): Spearman correlation coefficient ($\rho$). (c): Pearson correlation coefficient ($r$).

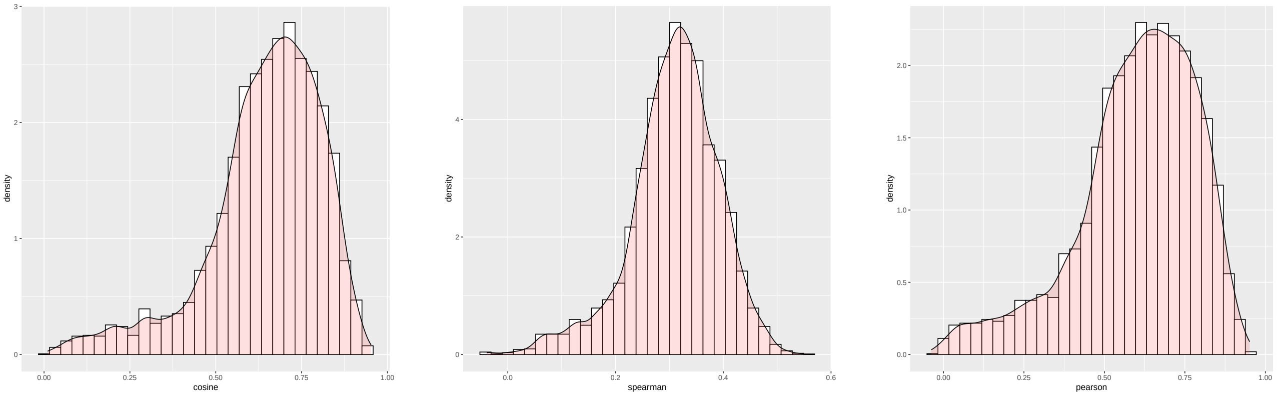


(a) (b) (c)

**Fig. 22.** The similarity scores of the sense and antisense counts of cells explored in section 2.5 for the mouse brain scMultiome dataset. (a): cosine similarity. (b): Spearman correlation coefficient ($\rho$). (c): Pearson correlation coefficient ($r$).

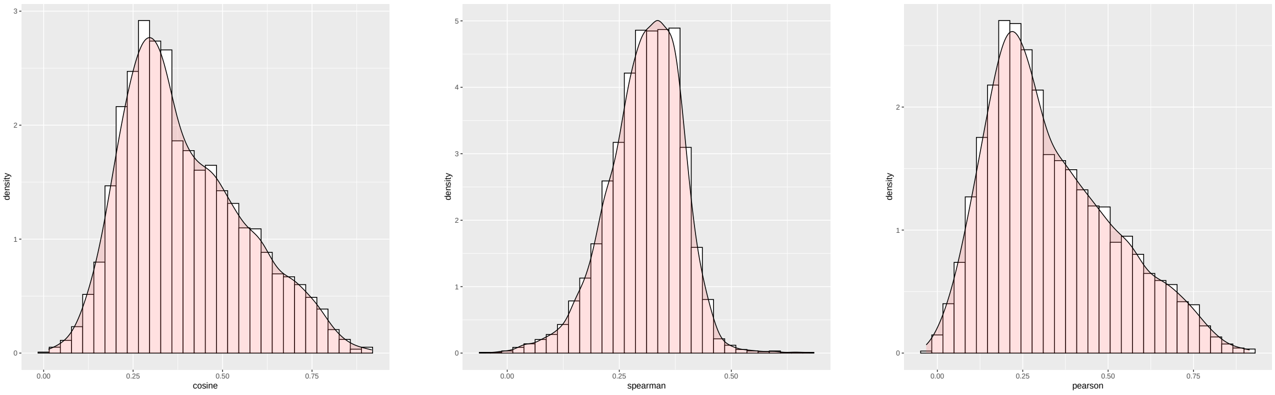


(a) (b) (c)

**Fig. 23.** The similarity scores of the sense and antisense counts of cells explored in section 2.5 for the mouse brain dataset. (a): cosine similarity. (b): Spearman correlation coefficient ($\rho$). (c): Pearson correlation coefficient ($r$).

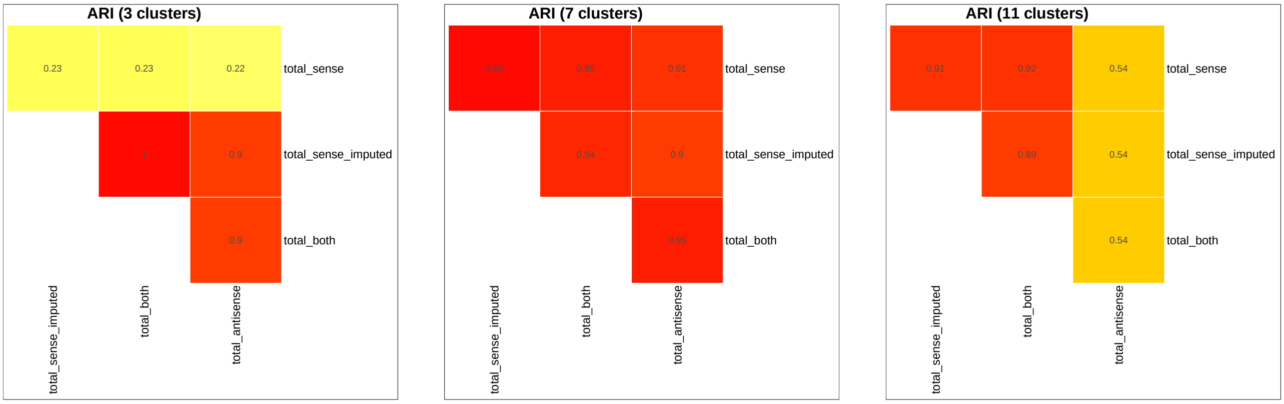


(a) (b) (c)

**Fig. 24.** The ARI of the clustering results generated from the standard and antisense modalities explored in section 2.5 for the human PBMC scMultiome dataset. `S`: spliced counts. `U`: unspliced counts. `A`: ambiguous counts. `SA`: spliced and ambiguous total counts. `USA`: spliced, unspliced, and ambiguous total counts. (a) represents the ARIs between the clusters discovered from different modalities using a low clustering resolution. (b) represents the results from a middle clustering resolution. (c) represents the results from a high clustering resolution.

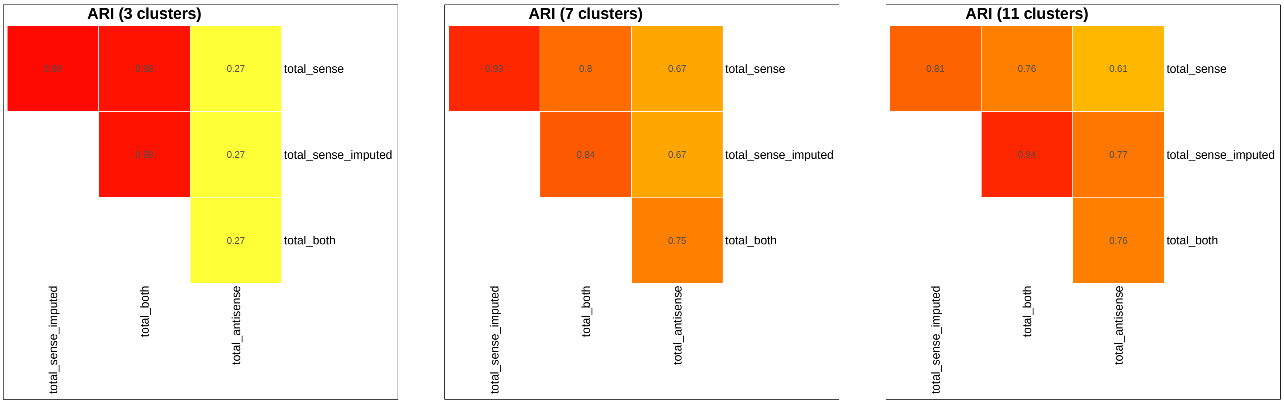


(a) (b) (c)

**Fig. 25.** The ARI of the clustering results generated from the standard and antisense modalities explored in section 2.5 for the human PBMC scRNA-seq dataset. `S`: spliced counts. `U`: unspliced counts. `A`: ambiguous counts. `SA`: spliced and ambiguous total counts. `USA`: spliced, unspliced and ambiguous total counts. (a) represents the ARIs between the clusters discovered from different modalities using a low clustering resolution. (b) represents the results from a middle clustering resolution. (c) represents the results from a high clustering resolution.

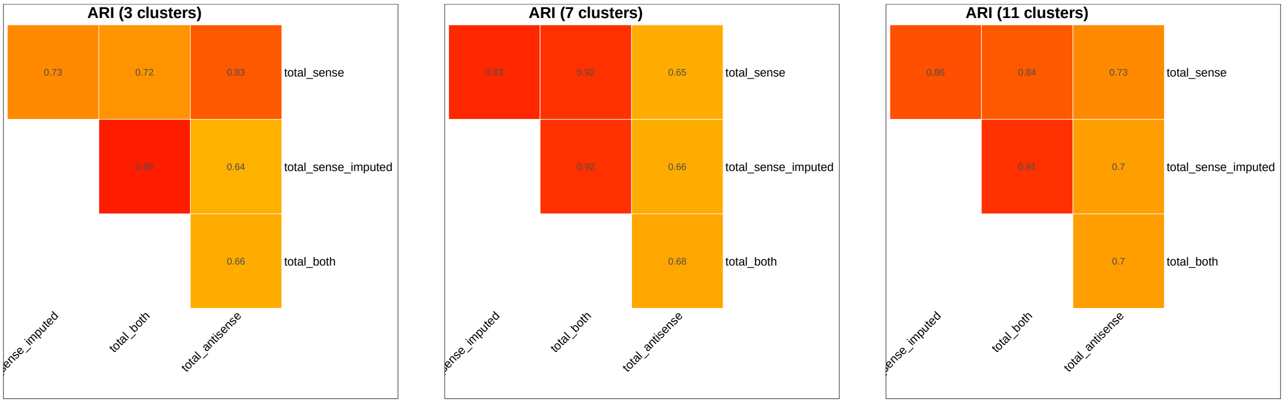


(a) (b) (c)

**Fig. 26.** The ARI of the clustering results generated from the standard and antisense modalities explored in section 2.5 for the human BMMC scMultiome dataset. `S`: spliced counts. `U`: unspliced counts. `A`: ambiguous counts. `SA`: spliced and ambiguous total counts. `USA`: spliced, unspliced, and ambiguous total counts. (a) represents the ARIs between the clusters discovered from different modalities using a low clustering resolution. (b) represents the results from a middle clustering resolution. (c) represents the results from a high clustering resolution.

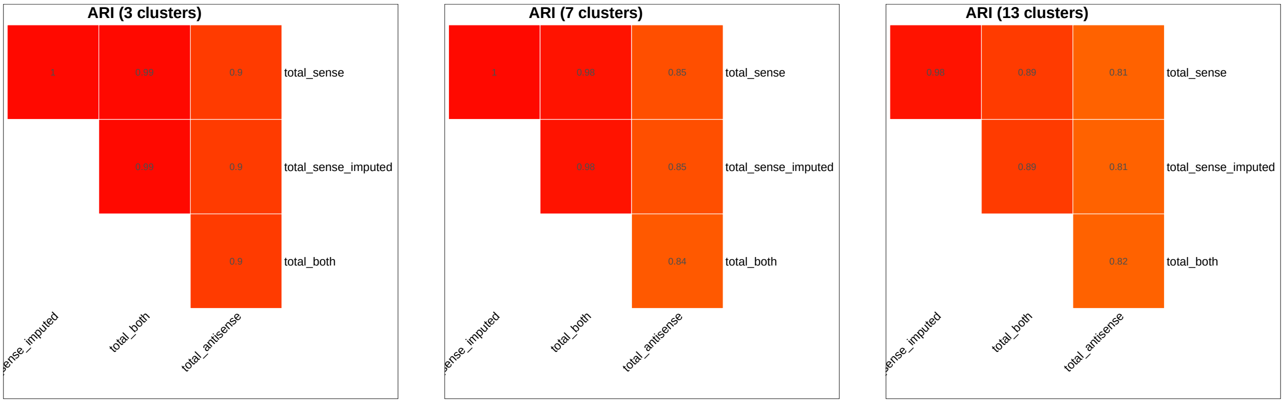


(a) (b) (c)

**Fig. 27.** The ARI of the clustering results generated from the standard and antisense modalities explored in section 2.5 for the human BMMC dataset. `S`: spliced counts. `U`: unspliced counts. `A`: ambiguous counts. `SA`: spliced and ambiguous total counts. `USA`: spliced, unspliced, and ambiguous total counts. (a) represents the ARIs between the clusters discovered from different modalities using a low clustering resolution. (b) represents the results from a middle clustering resolution. (c) represents the results from a high clustering resolution.

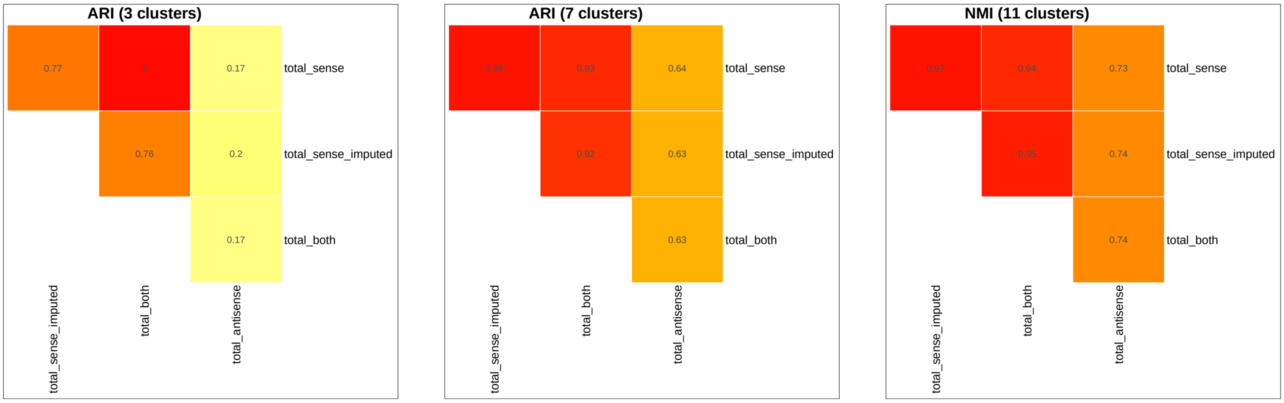


(a) (b) (c)

**Fig. 28.** The ARI of the clustering results generated from the standard and antisense modalities explored in section 2.5 for the human brain scMultiome dataset. `S`: spliced counts. `U`: unspliced counts. `A`: ambiguous counts. `SA`: spliced and ambiguous total counts. `USA`: spliced, unspliced, and ambiguous total counts. (a) represents the ARIs between the clusters discovered from different modalities using a low clustering resolution. (b) represents the results from a middle clustering resolution. (c) represents the results from a high clustering resolution.

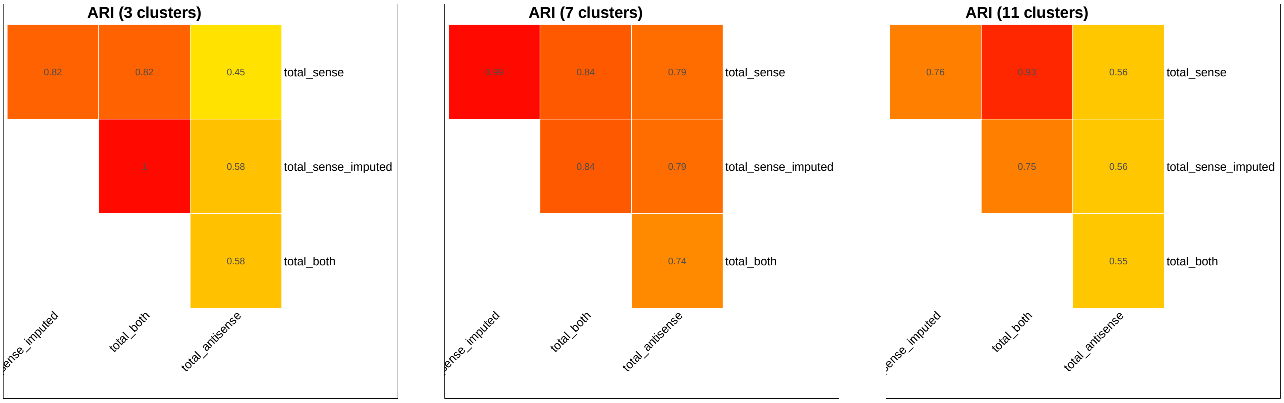


(a) (b) (c)

**Fig. 29.** The ARI of the clustering results generated from the standard and antisense modalities explored in section 2.5 for the mouse brain scMultiome dataset. `S`: spliced counts. `U`: unspliced counts. `A`: ambiguous counts. `SA`: spliced and ambiguous total counts. `USA`: spliced, unspliced, and ambiguous total counts. (a) represents the ARIs between the clusters discovered from different modalities using a low clustering resolution. (b) represents the results from a middle clustering resolution. (c) represents the results from a high clustering resolution.

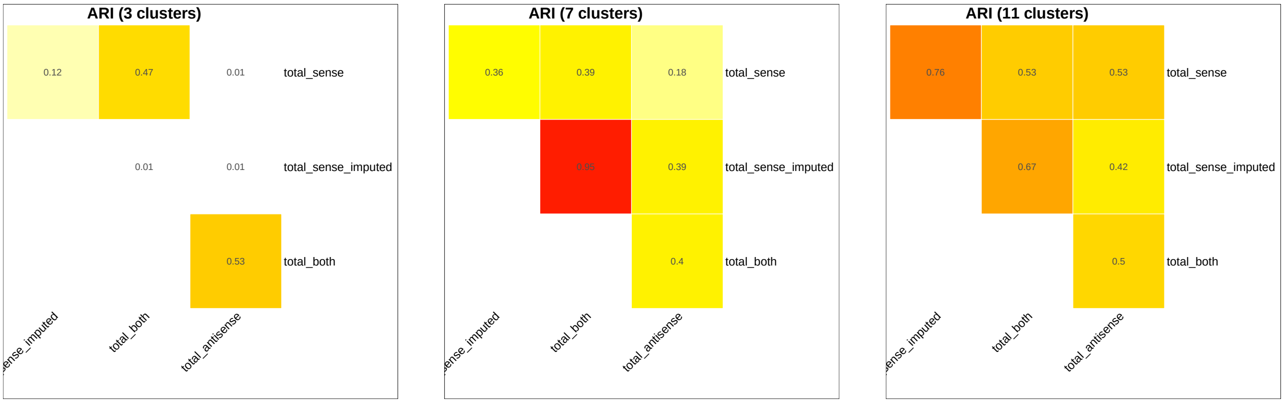


(a) (b) (c)

**Fig. 30.** The ARI of the clustering results generated from the standard and antisense modalities explored in section 2.5 for the mouse brain dataset. `S`: spliced counts. `U`: unspliced counts. `A`: ambiguous counts. `SA`: spliced and ambiguous total counts. `USA`: spliced, unspliced, and ambiguous total counts. (a) represents the ARIs between the clusters discovered from different modalities using a low clustering resolution. (b) represents the results from a middle clustering resolution. (c) represents the results from a high clustering resolution.

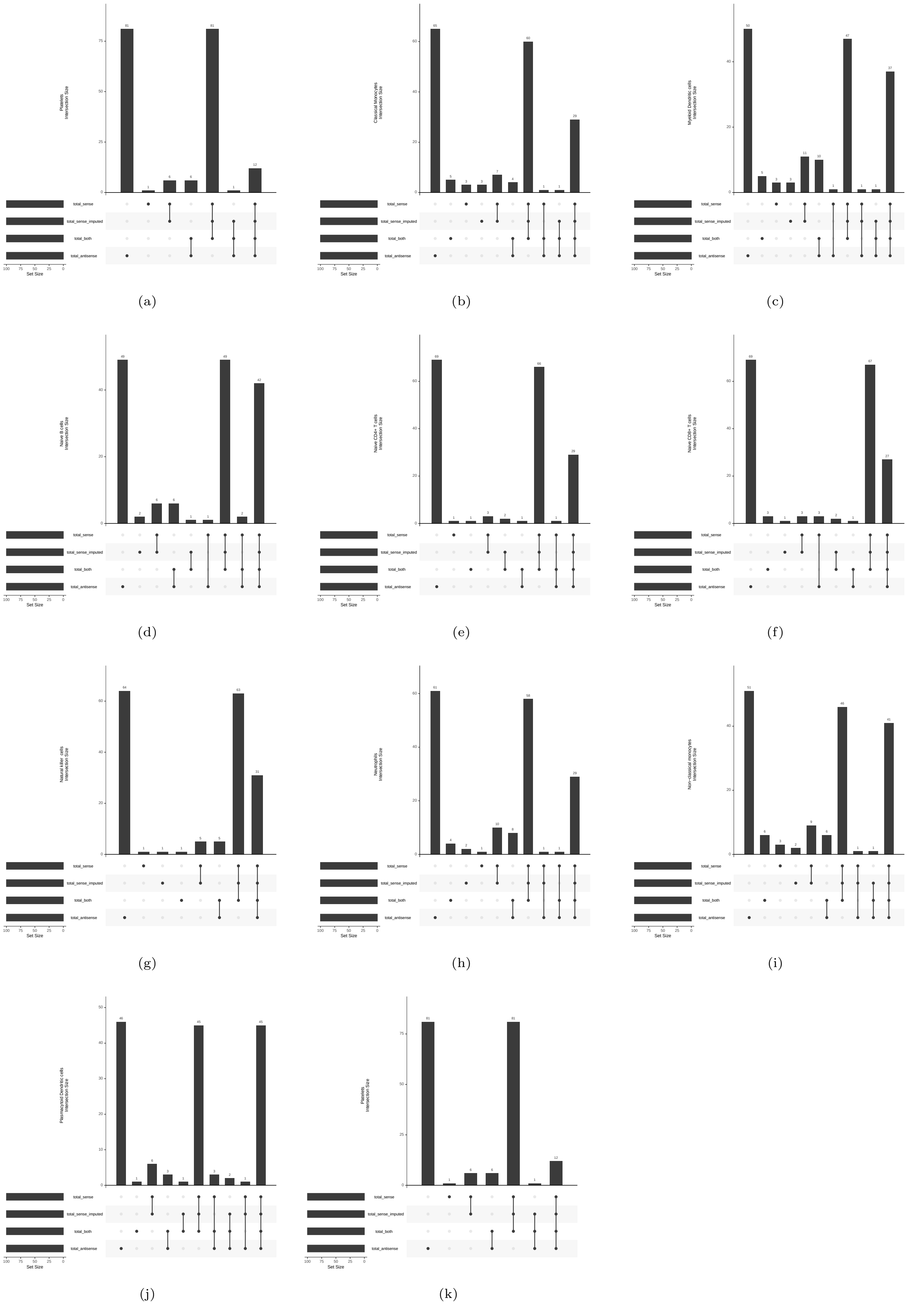


**Fig. 31.** The differentially expressed genes of each `sctype` cell type discovered from the antisense transcriptomic count matrices of the human PBMC scRNA-seq dataset. Cell types are listed as a part of the y-axis label of the histogram.

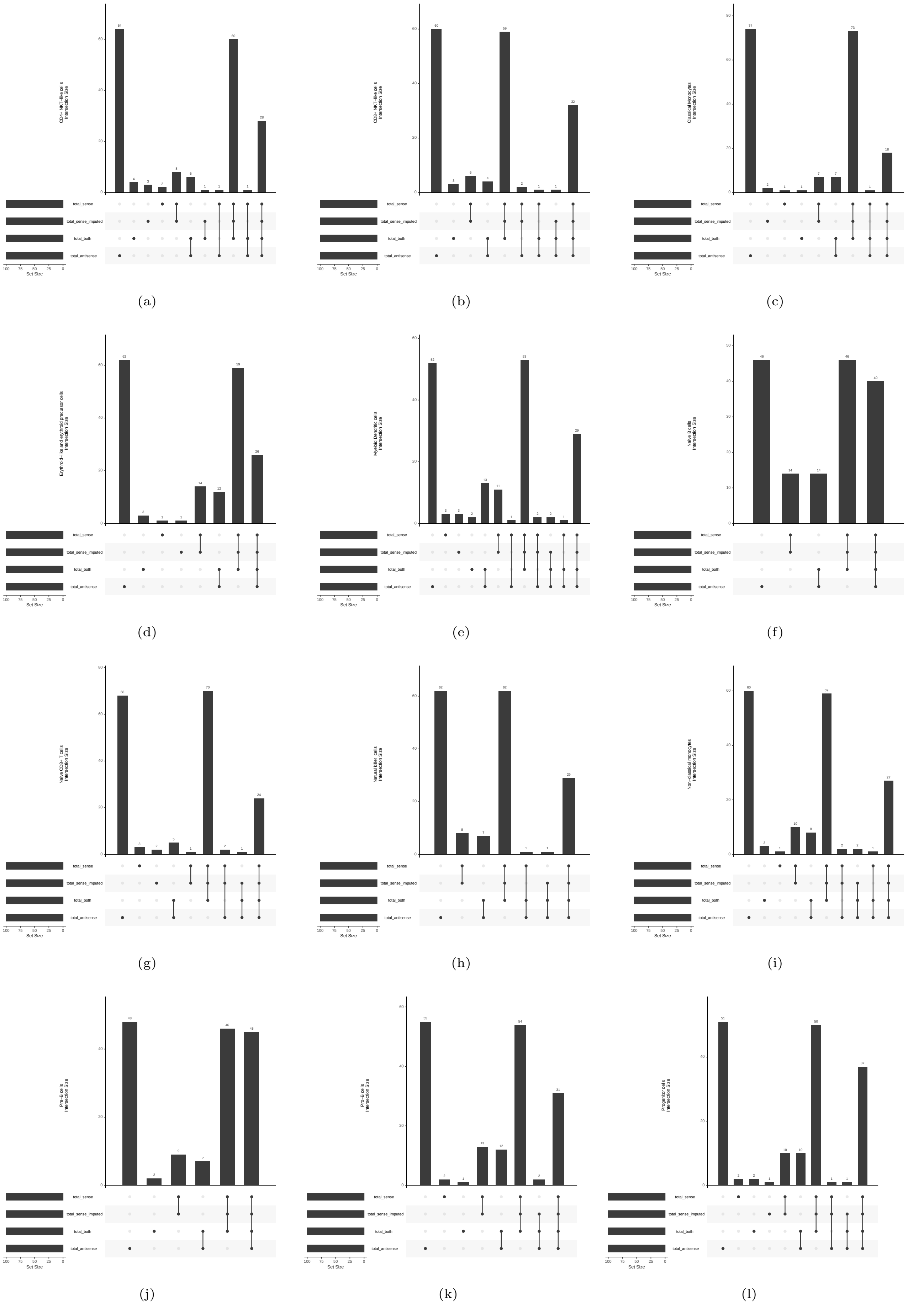


**Fig. 32.** The differentially expressed genes of each `sctype` cell type discovered from the antisense transcriptomic count matrices of the human BMMC scRNA-seq dataset. Cell types are listed as a part of the y-axis label of the histogram.

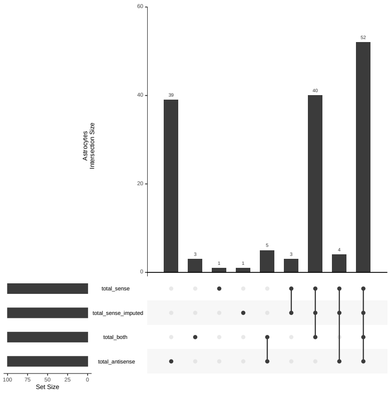


(a)

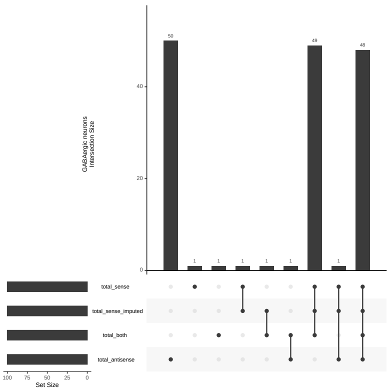


(b)

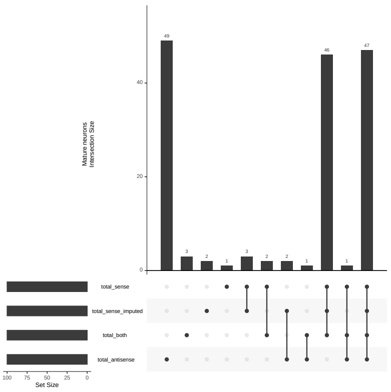


(c)

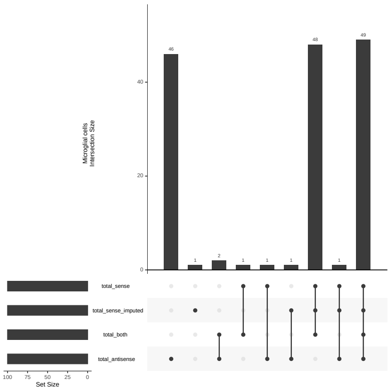


(d)

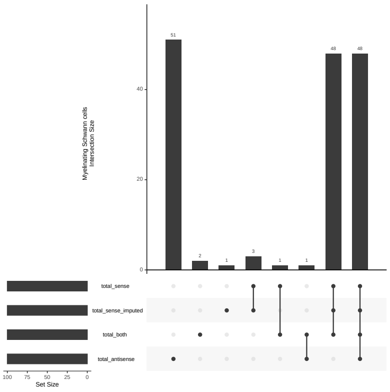


(e)

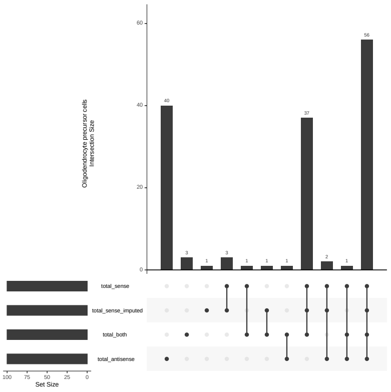


(f)

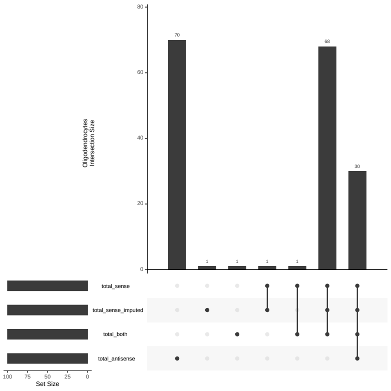


(g)

**Fig. 33.** The differentially expressed genes of each `sctype` cell type discovered from the antisense transcriptomic count matrices of the human brain scRNA-seq dataset. Cell types are listed as a part of the y-axis label of the histogram. The results are from the *scRNA-seq optimized gene annotations*.

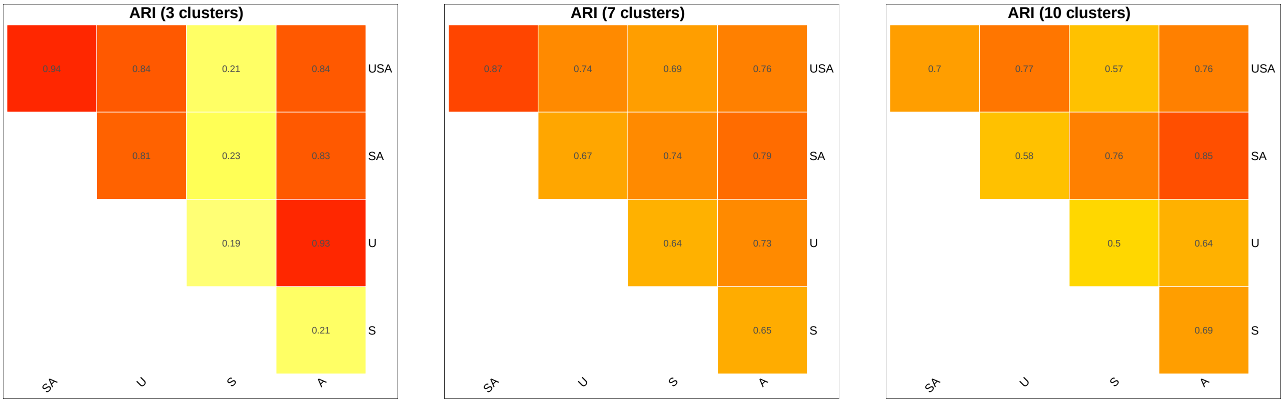


(a) (b) (c)

**Fig. 34.** The ARI of the clustering results generated from the five sense transcriptomic modalities explored in section 2.4 for the human PBMC scMultiome scRNA-seq dataset. `S`: spliced counts. `U`: unspliced counts. `A`: ambiguous counts. `SA`: spliced and ambiguous total counts. `USA`: spliced, unspliced, and ambiguous total counts. (a) represents the ARIs between the clusters discovered from different modalities using a low clustering resolution. (b) represents the results from a middle clustering resolution. (c) represents the results from a high clustering resolution.

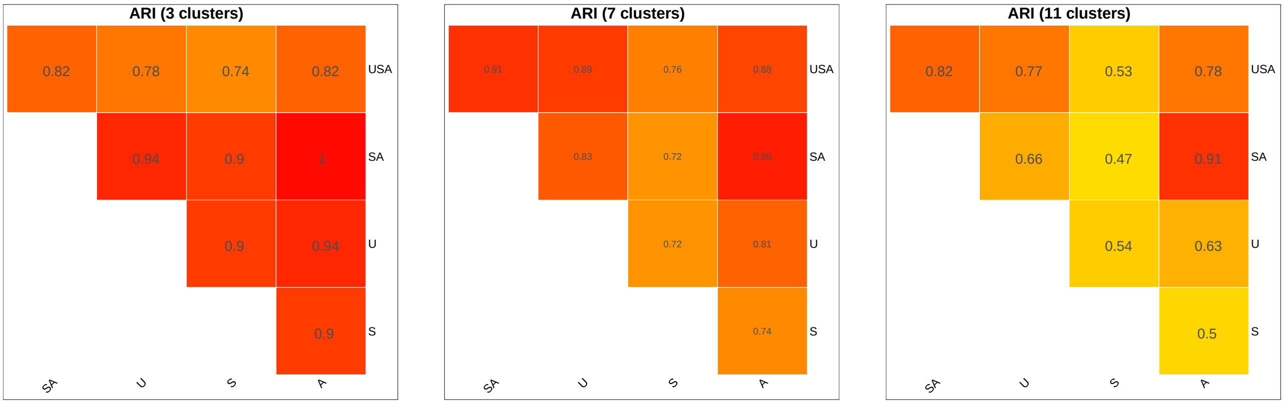


(a) (b) (c)

**Fig. 35.** The ARI of the clustering results generated from the five sense transcriptomic modalities explored in section 2.4 for the human PBMC scRNA-seq dataset. `S`: spliced counts. `U`: unspliced counts. `A`: ambiguous counts. `SA`: spliced and ambiguous total counts. `USA`: spliced, unspliced, and ambiguous total counts. (a) represents the ARIs between the clusters discovered from different modalities using a low clustering resolution. (b) represents the results from a middle clustering resolution. (c) represents the results from a high clustering resolution.

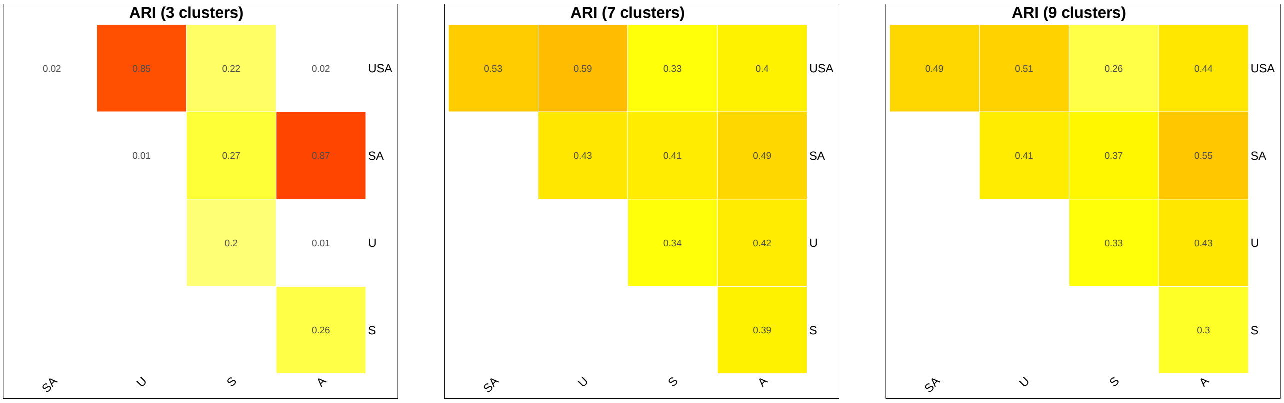


(a) (b) (c)

**Fig. 36.** The ARI of the clustering results generated from the five sense transcriptomic modalities explored in section 2.4 for the human BMMC scMultiome dataset. `S`: spliced counts. `U`: unspliced counts. `A`: ambiguous counts. `SA`: spliced and ambiguous total counts. `USA`: spliced, unspliced, and ambiguous total counts. (a) represents the ARIs between the clusters discovered from different modalities using a low clustering resolution. (b) represents the results from a middle clustering resolution. (c) represents the results from a high clustering resolution.

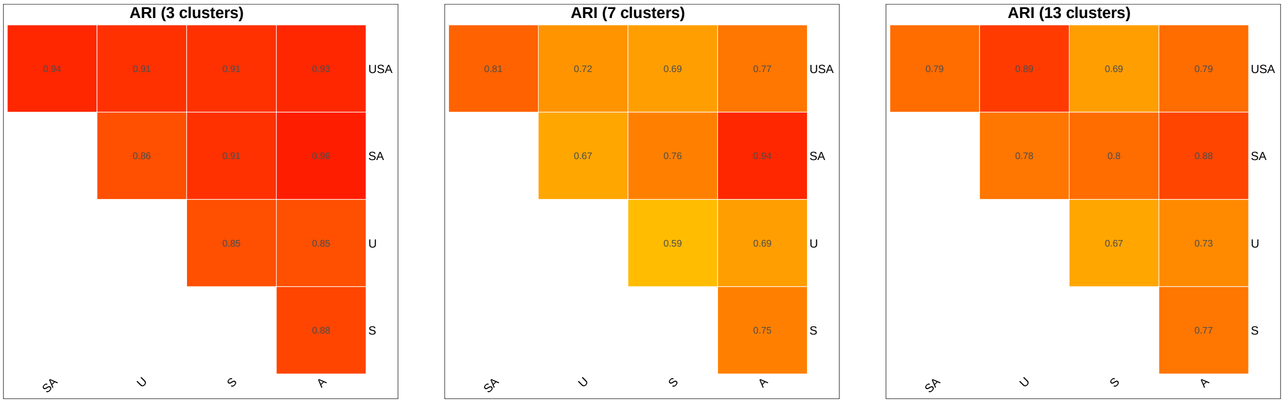


(a) (b) (c)

**Fig. 37.** The ARI of the clustering results generated from the five sense transcriptomic modalities explored in section 2.4 for the human BMMC dataset. `S`: spliced counts. `U`: unspliced counts. `A`: ambiguous counts. `SA`: spliced and ambiguous total counts. `USA`: spliced, unspliced, and ambiguous total counts. (a) represents the ARIs between the clusters discovered from different modalities using a low clustering resolution. (b) represents the results from a middle clustering resolution. (c) represents the results from a high clustering resolution.

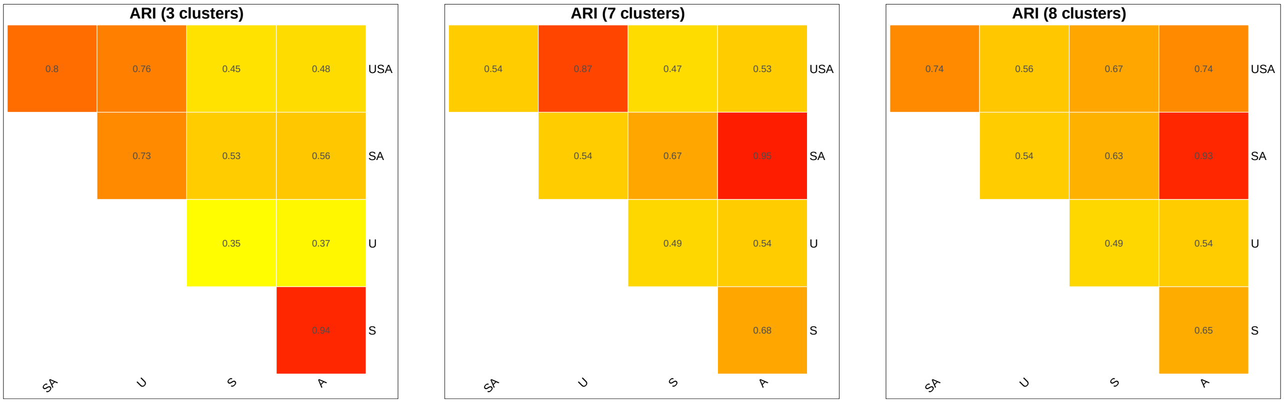


(a) (b) (c)

**Fig. 38.** The ARI of the clustering results generated from the five sense transcriptomic modalities explored in section 2.4 for the human brain scMultiome dataset. `S`: spliced counts. `U`: unspliced counts. `A`: ambiguous counts. `SA`: spliced and ambiguous total counts. `USA`: spliced, unspliced, and ambiguous total counts. (a) represents the ARIs between the clusters discovered from different modalities using a low clustering resolution. (b) represents the results from a middle clustering resolution. (c) represents the results from a high clustering resolution.

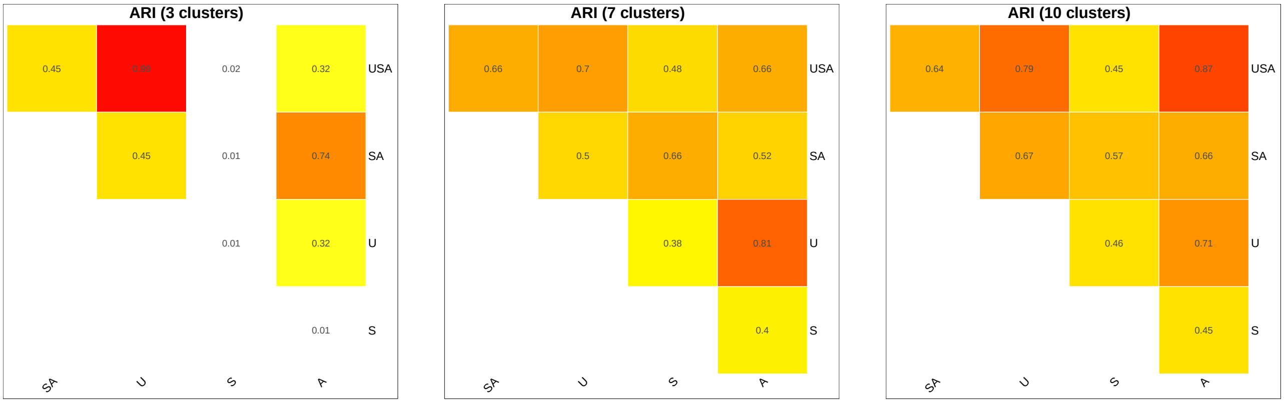


(a) (b) (c)

**Fig. 39.** The ARI of the clustering results generated from the five sense transcriptomic modalities explored in section 2.4 for the mouse brain scMultiome dataset. `S`: spliced counts. `U`: unspliced counts. `A`: ambiguous counts. `SA`: spliced and ambiguous total counts. `USA`: spliced, unspliced, and ambiguous total counts. (a) represents the ARIs between the clusters discovered from different modalities using a low clustering resolution. (b) represents the results from a middle clustering resolution. (c) represents the results from a high clustering resolution.

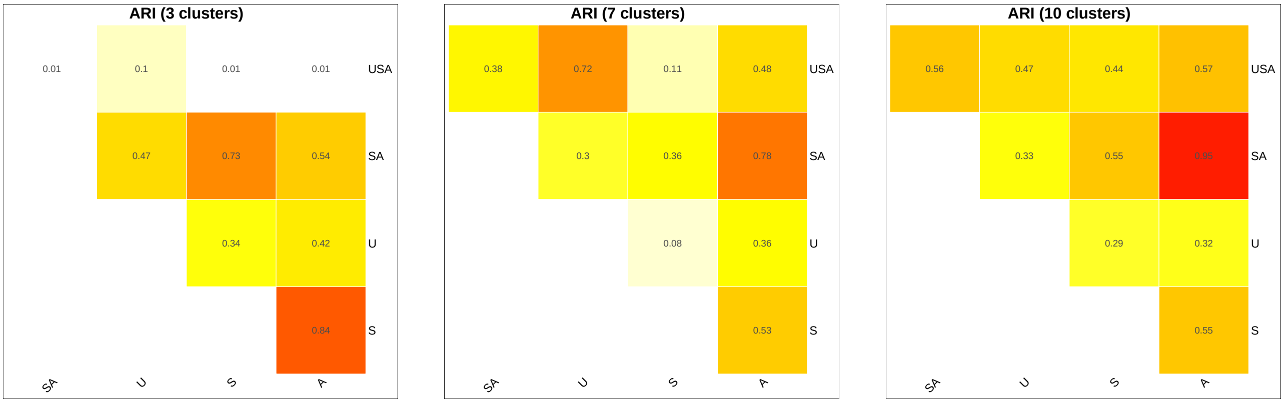


(a) (b) (c)

**Fig. 40.** The ARI of the clustering results generated from the five sense transcriptomic modalities explored in section 2.4 for the mouse brain dataset. `S`: spliced counts. `U`: unspliced counts. `A`: ambiguous counts. `SA`: spliced and ambiguous total counts. `USA`: spliced, unspliced, and ambiguous total counts. (a) represents the ARIs between the clusters discovered from different modalities using a low clustering resolution. (b) represents the results from a middle clustering resolution. (c) represents the results from a high clustering resolution.

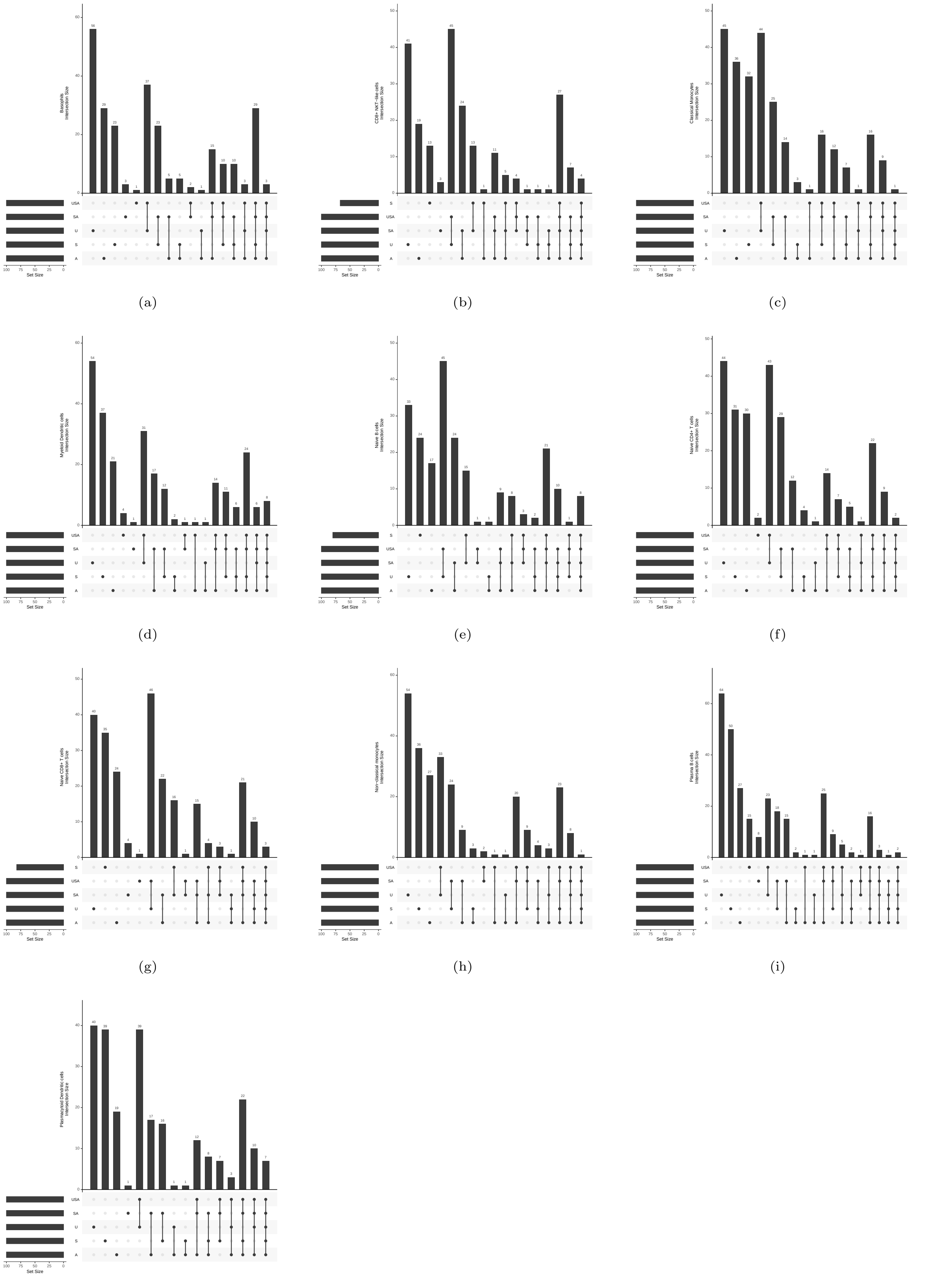


(a) (b) (c)

(d) (e) (f)

(g) (h) (i)

(j)

**Fig. 41.** The differentially expressed genes of each sctype cell type discovered from the sense transcriptomic count matrices of the human PBMC scMultiome scRNA-seq dataset. Cell types are listed as a part of the y-axis label of the histogram.

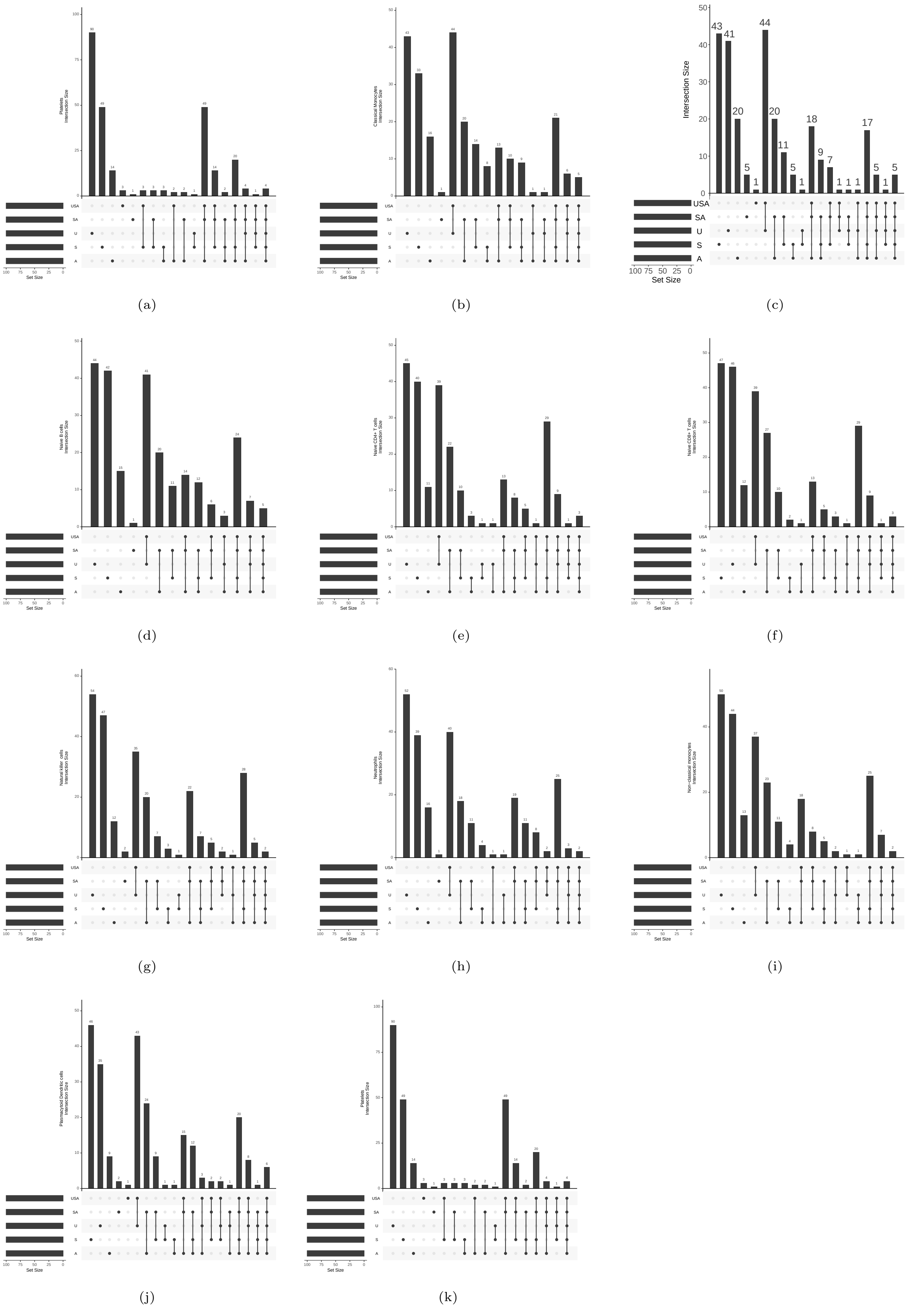


**Fig. 42.** The differentially expressed genes of each `sctype` cell type discovered from the sense transcriptomic count matrices of the human PBMC scRNA-seq dataset. Cell types are listed as a part of the y-axis label of the histogram.

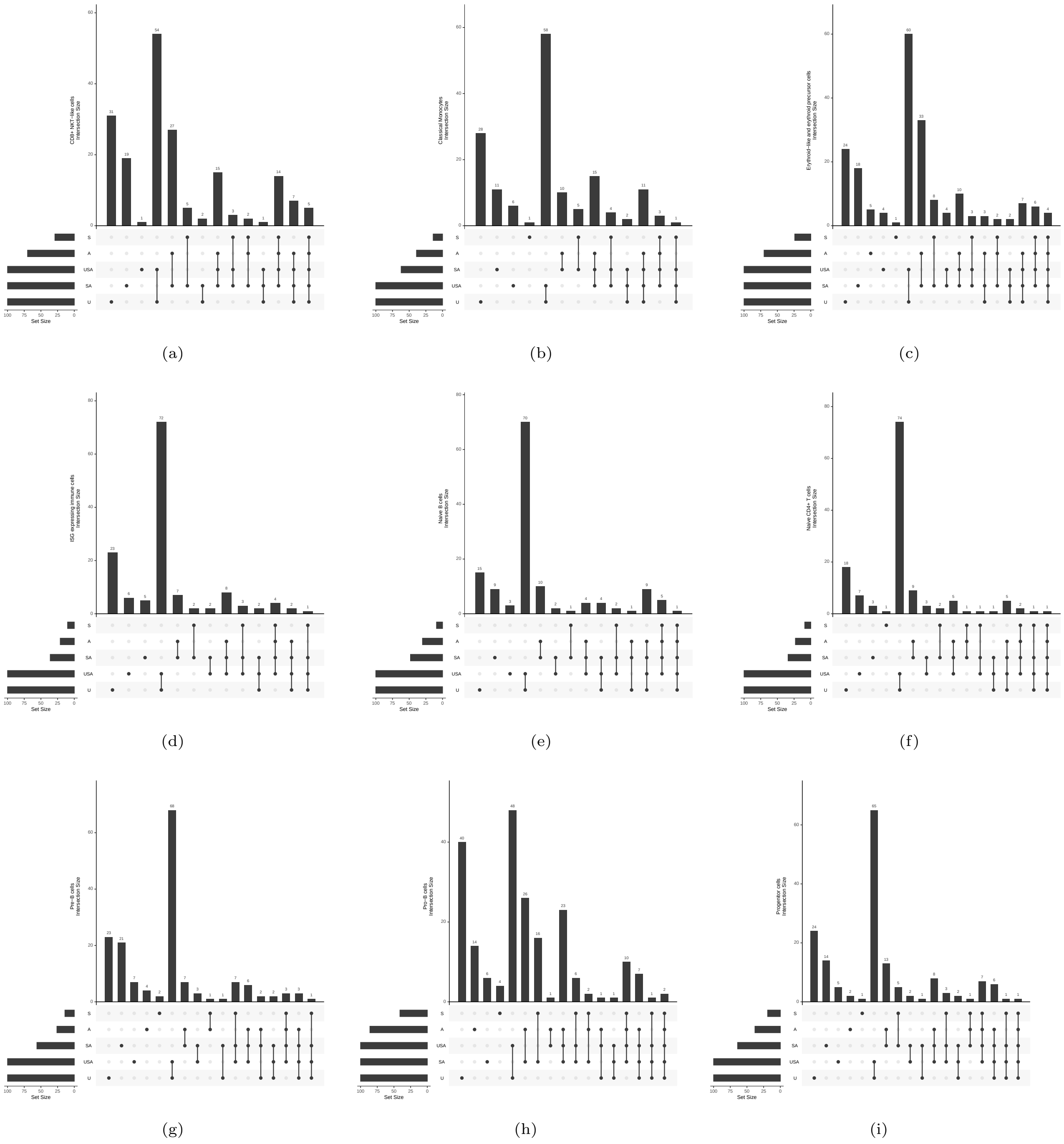


**Fig. 43.** The differentially expressed genes of each `sctype` cell type discovered from the sense transcriptomic count matrices of the human BMMC scMultiome dataset. Cell types are listed as a part of the y-axis label of the histogram.

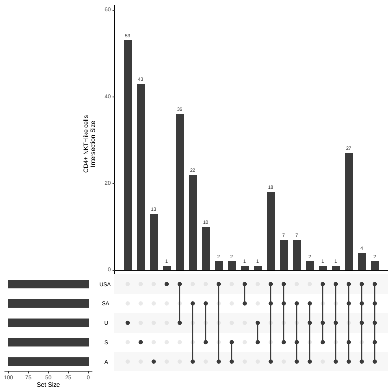


(a)

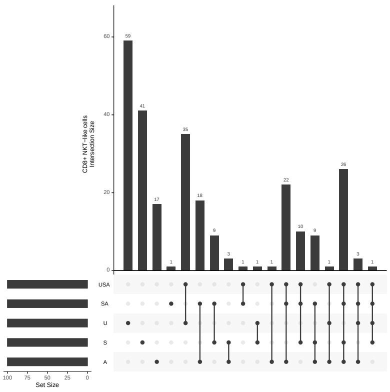


(b)

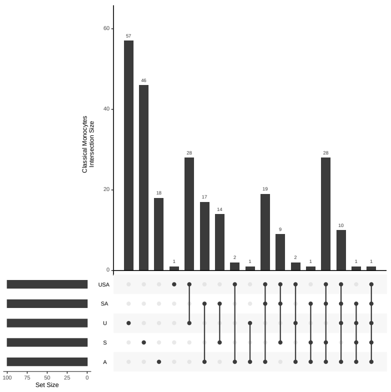


(c)

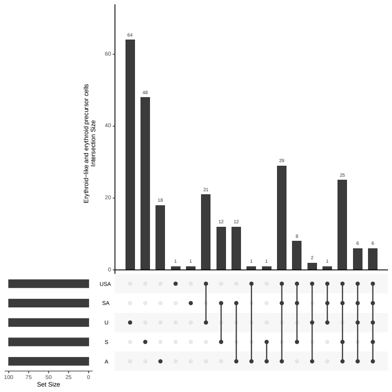


(d)

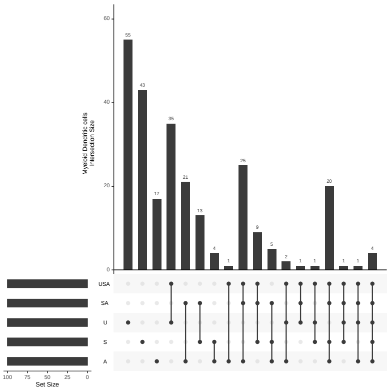


(e)

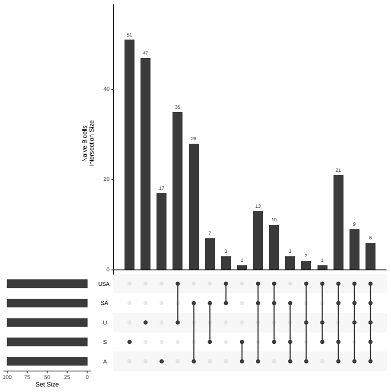


(f)

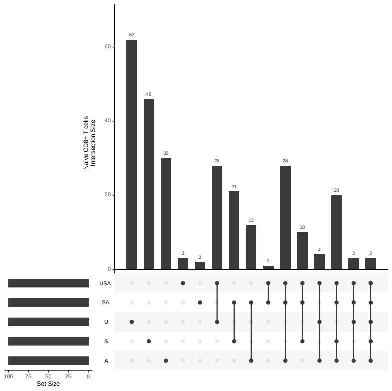


(g)

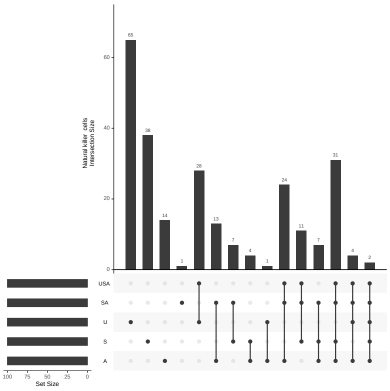


(h)

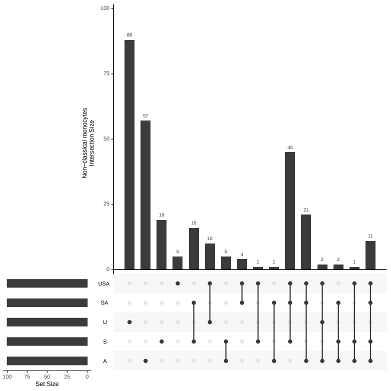


(i)

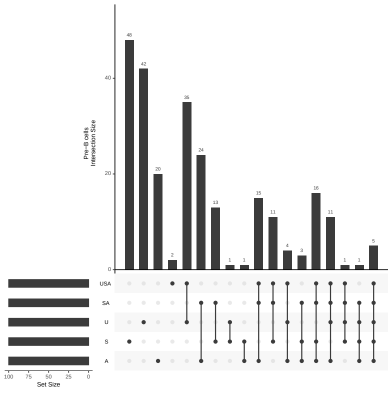


(j)

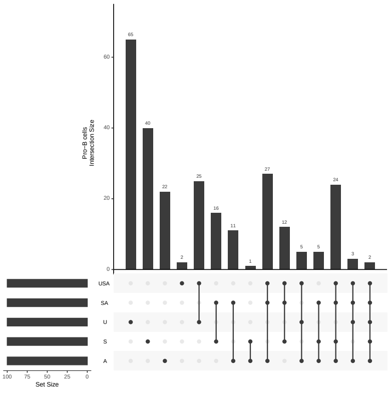


(k)

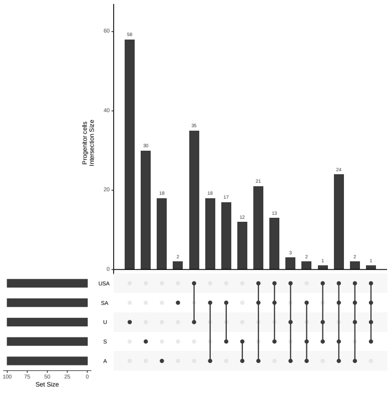


(l)

**Fig. 44.** The differentially expressed genes of each `sctype` cell type discovered from the sense transcriptomic count matrices of the human BMMC scRNA-seq dataset. Cell types are listed as a part of the y-axis label of the histogram.

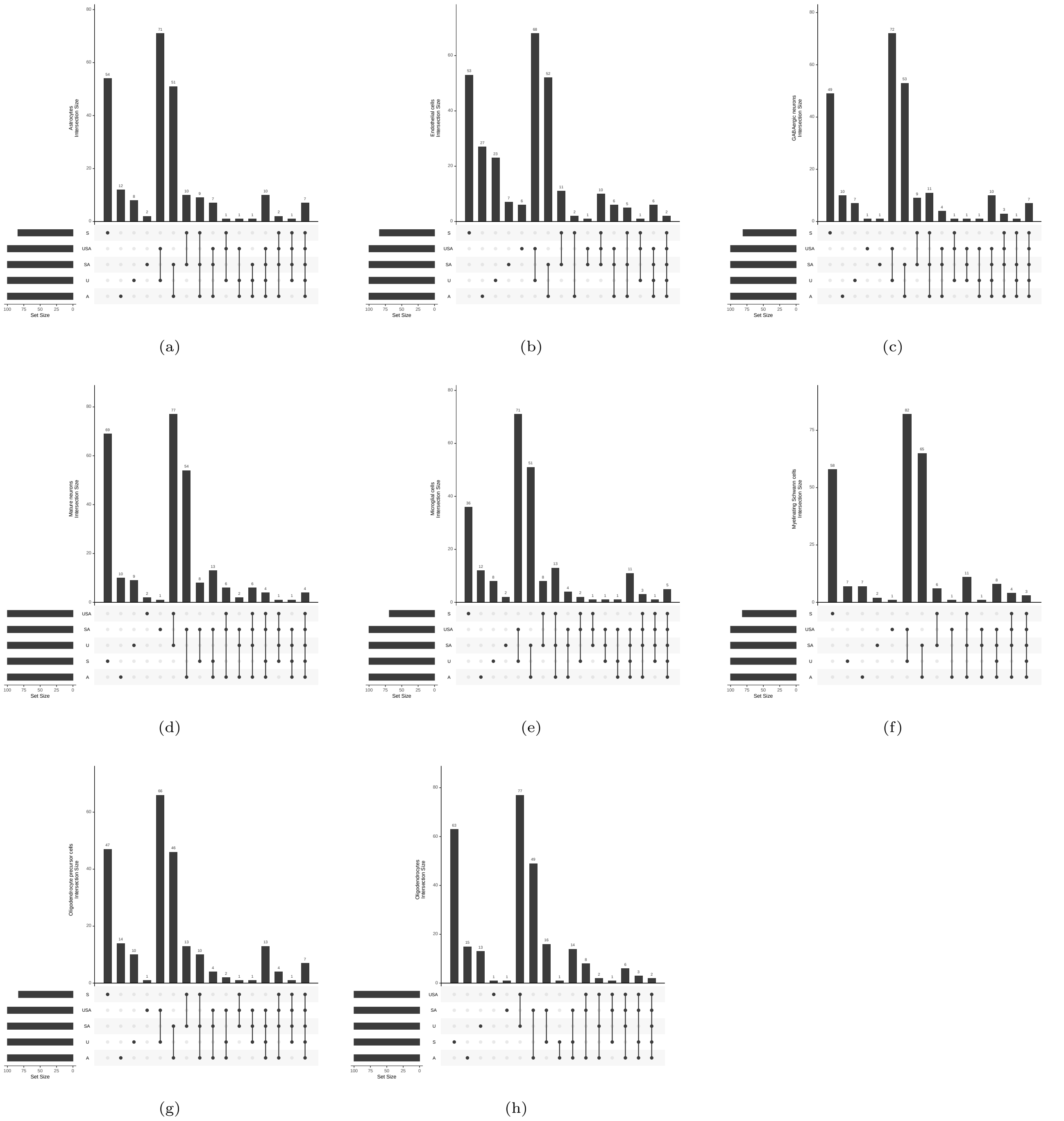


**Fig. 45.** The differentially expressed genes of each `sctype` cell type discovered from the sense transcriptomic count matrices of the human brain scMultiome dataset. Cell types are listed as a part of the y-axis label of the histogram.

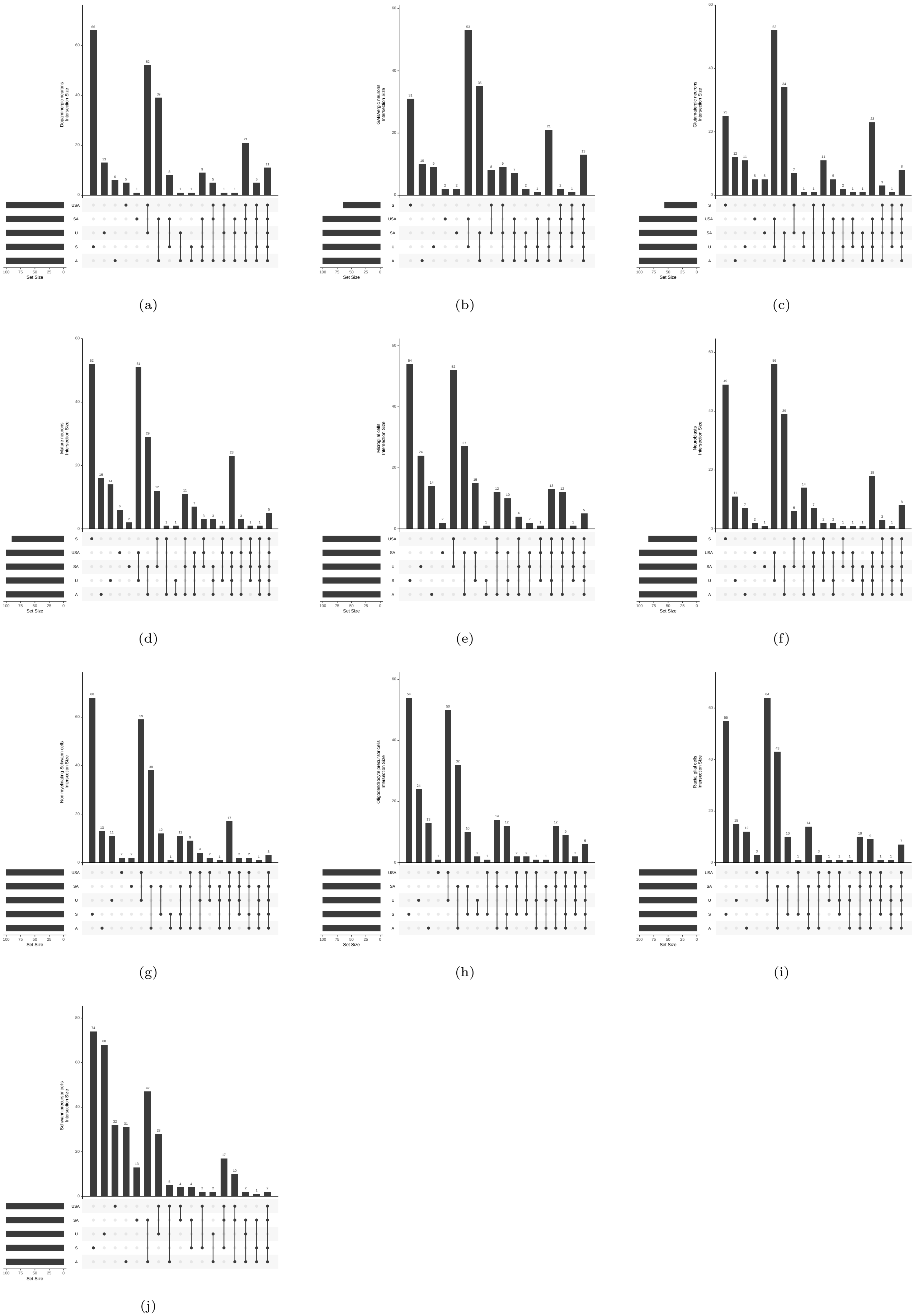


**Fig. 46.** The differentially expressed genes of each sctype cell type discovered from the sense transcriptomic count matrices of the mouse brain scMultiome dataset. Cell types are listed as a part of the y-axis label of the histogram.

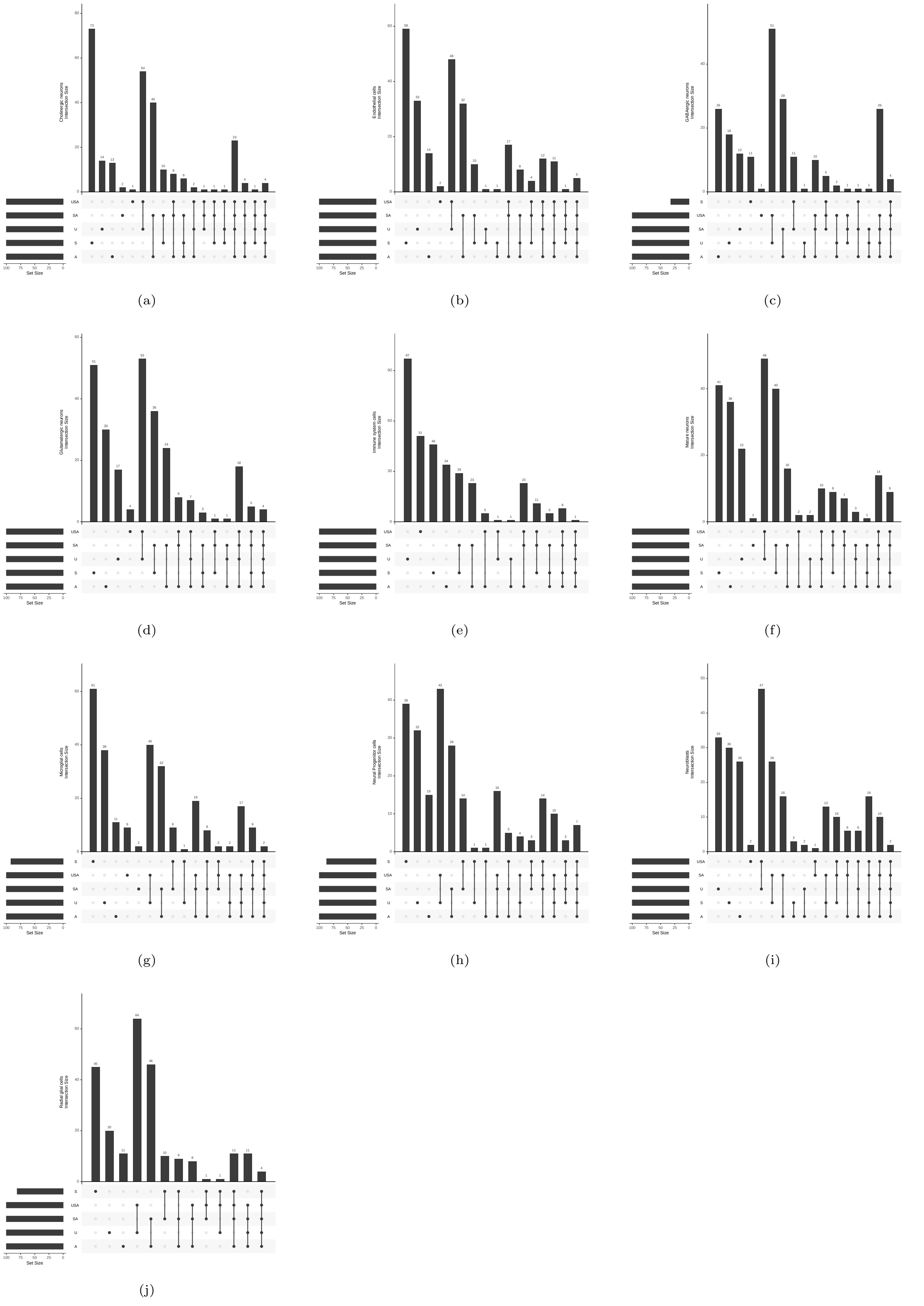


**Fig. 47.** The differentially expressed genes of each sctype cell type discovered from the sense transcriptomic count matrices of the mouse brain scRNA-seq dataset. Cell types are listed as a part of the y-axis label of the histogram.
